# Reduced hybrid survival in a migratory divide between songbirds

**DOI:** 10.1101/2024.01.23.576948

**Authors:** Stephanie A. Blain, Hannah C. Justen, Wendy Easton, Kira E. Delmore

## Abstract

Migratory divides, hybrid zones between populations that use different seasonal migration routes, are hypothesized to contribute to speciation. Specifically, relative to parental species, hybrids at divides are predicted to exhibit (1) intermediate migratory behavior and (2) reduced fitness as a result. We provide the first direct test of the second prediction here with one of the largest existing avian tracking datasets, leveraging a divide between Swainson’s thrushes where the first prediction is supported. Using detection rates as a proxy for survival, our results supported the migratory divide hypothesis with lower survival rates for hybrids than parental forms. This finding was juvenile-specific (vs. adults), suggesting selection against hybrids is stronger earlier in life. Reduced hybrid survival was not explained by selection against intermediate phenotypes or negative interactions among phenotypes. Additional work connecting specific features of migration is needed, but these patterns provide strong support for migration as an ecological driver of speciation.

## Introduction

Divergent ecological selection can contribute to speciation in several ways, including extrinsic postzygotic isolation, where hybrids are poorly-adapted to parental niches (Dieckmann and Doebeli 1999; Schluter 2000; Dettman et al. 2007). Robust theoretical support for ecological speciation exists but few well-supported examples of extrinsic postzygotic isolation have been described in nature (Benkman 2003; Arnegard et al. 2014). Migratory divides, hybrid zones between closely related populations that use different routes to travel between breeding and nonbreeding sites (Irwin and Irwin 2005; Møller et al. 2011; Rohwer and Irwin 2011), may help fill this empirical gap in the literature. Migratory routes and associated differences in migratory phenotypes (e.g., orientation, timing and morphology) are genetically determined in many groups and thus it has been predicted that (1) hybrids will exhibit intermediate migratory phenotypes and (2) as a result, hybrids will fall between parental niches and exhibit reduced fitness relative to parental forms (Helbig 1991*a*; Irwin and Irwin 2005; Turbek et al. 2017). Support for the first prediction from this hypothesis (hereafter the ‘migratory divide hypothesis’) has been demonstrated, with hybrids exhibiting intermediate migratory routes (Delmore and Irwin 2014; Vali et al. 2018; Sokolovskis et al. 2023). Whether these routes reduce the fitness of hybrids (i.e., the second prediction) has yet to be tested directly.

There are two mechanisms by which hybrid fitness can be reduced in migratory divides. First, parental populations in divides often circumvent large geographic barriers with inhospitable habitat, including the Tibetan Plateau in barn swallows (Scordato et al. 2020), the Rocky Mountains in monarch butterflies (Reppert et al. 2010), and the northern Atlantic Ocean in eels (Albert et al. 2006). If migratory behaviour is additively inherited, hybrids could have intermediate trajectories, stopover sites, or nonbreeding locations that bring them over these barriers, leading to poor performance on migration and ultimately low survival or fecundity (Irwin and Irwin 2005; Newton 2006). Second, mismatch may also contribute to reductions in fitness. Specifically, migration is also a syndrome that requires the integration of many phenotypes (behavioural, morphological, and physiological; Dingle 2006; Justen and Delmore 2022). Accordingly, ecological selection may instead act on sets of phenotypes that are not coordinated well with each other (i.e., are mismatched; Arnegard et al. 2014; Chhina et al. 2022). For example, a hybrid individual may have the migratory timing of one species and the wing shape of the other. Migration would act as an extrinsic postzygotic barrier to gene flow under both scenarios (intermediate routes or mismatched migratory phenotypes).

Conclusive evidence for extrinsic postzygotic isolation requires direct estimates of ecological selection against hybrids in nature (Rundle and Nosil 2005). In the case of the migratory divide hypothesis, this would require individual-level survival data from migration. Advances in animal movement ecology have allowed researchers to start tracking individual birds over the entire annual cycle (Bridge et al. 2011). However, most technologies (e.g., archival tags) store location information in the tag and therefore require recapture, making it impossible to know the migration phenotype of the birds that did not survive. Additionally, archival tags are only useful for adult birds because juvenile (hatch-year) birds typically disperse from their natal grounds (Paradis et al. 1998), making recapture infeasible. Selection on juveniles may be more relevant than selection on adults, as adults have already survived migration at least one year and mortality is generally higher during their first year of migration (Sergio et al. 2019; Robinson et al. 2020). Automated radio telemetry systems, such as the globally distributed Motus station network (Fig. 1A), provide an exciting solution to this problem as birds do not need to be recaptured; their locations are recorded when they pass by stations on migration (Taylor et al. 2017), making it possible to obtain estimates of survival and migratory behaviour for juveniles during migration.

**Fig. 1.**
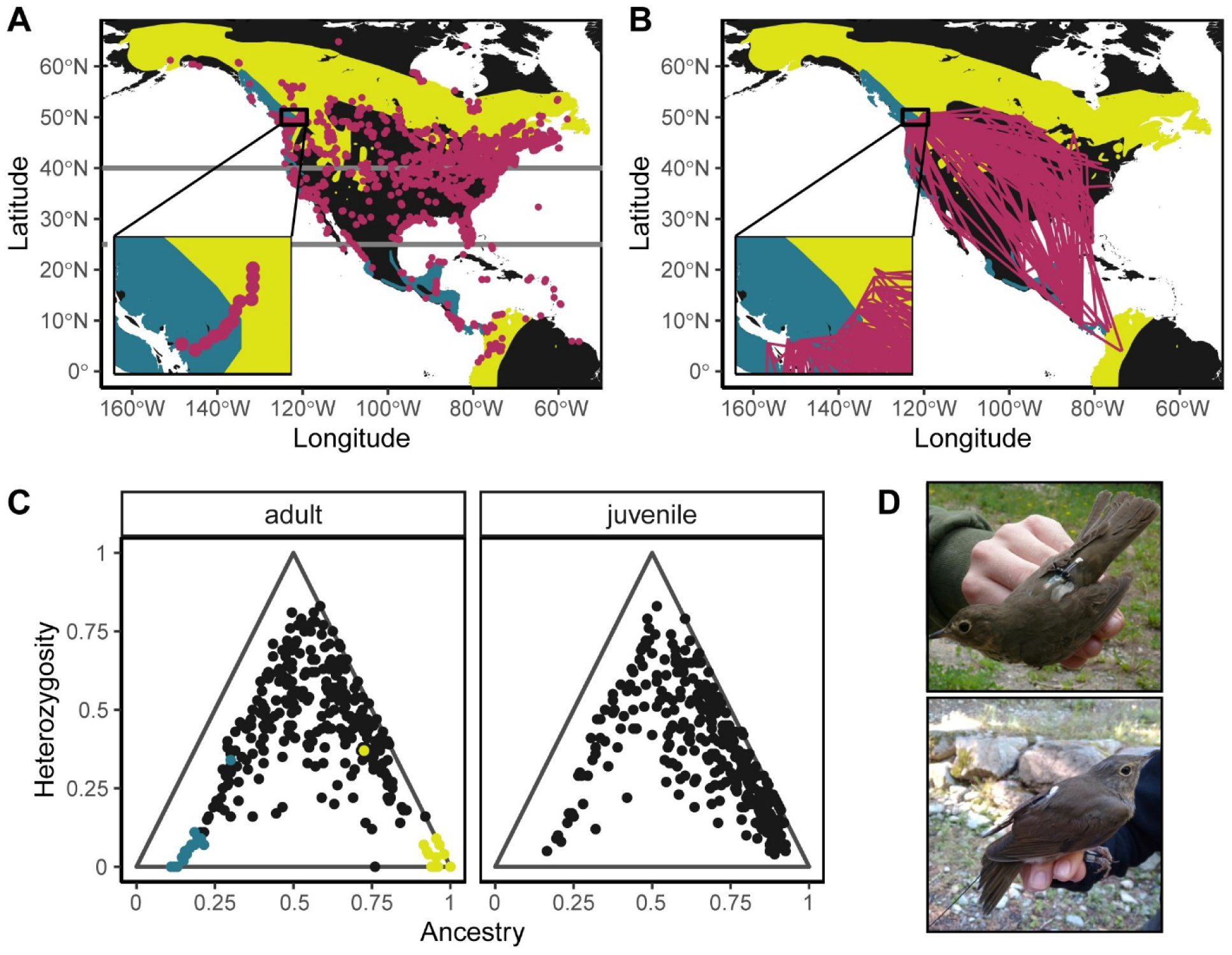
Geographic distribution, migratory tracks, and hybrid zone composition. (A) Breeding and nonbreeding ranges for inland (yellow) and coastal (blue) subspecies of Swainson’s thrushes. Points show locations of Motus receiver stations active at some time between August 2019 – 2023. Horizontal grey lines indicate the latitude bands used as cut-offs for binning survival into stages. Inset shows the fence of Motus receiver stations that span the hybrid zone in the southwestern British Columbia. (B) Radio station detections on fall migration for juvenile birds. Lines connect detections across stations for each individual. Inset shows tracks of juvenile birds detected while flying past the fence of stations in southwestern British Columbia. (C) Genomic ancestry and heterozygosity for adult and juvenile birds. Each point is one individual, with colour indicating whether birds were captured in the hybrid zone (grey), coastal subspecies range (blue), or inland subspecies range (yellow).(D) Image of an adult Swainson’s thrush with an archival tag (top) and a juvenile Swainson’s thrush with a radio tag (bottom).

Two Swainson’s thrush subspecies form a well-characterized migratory divide in western North America. The hybrid zone between these subspecies is narrow with low population densities at the centre (Ruegg 2008), suggesting that it is maintained by selection against hybrids (Barton and Hewitt 1985). Direct tracking data and ecological models suggest that differences in migration are responsible for poor hybrid fitness (Delmore and Irwin 2014; Justen et al. 2021).

Specifically, the subspecies take different routes on migration (Delmore et al. 2012). The coastal subspecies follows a migration route west of the Rocky Mountains, along the Pacific Coast, and overwinters in southern Mexico and Central America. The inland subspecies migrates east of the Rocky Mountains, over central and eastern North America, to northern South America. Hybrids exhibit considerable variation in the routes they take, including intermediate routes (Delmore and Irwin 2014). Ecological models suggested that these hybrid routes are inferior to parental routes; landscape connectivity, which reflects the level of movement permitted by landscape features, and habitat suitability at stopover sites were both lower along intermediate routes taken by hybrids relative to those of parental forms (Justen et al. 2021). However, a direct test of individual-level survival over the course of migration is needed to determine if hybrids exhibit reduced fitness relative to parental subspecies on migration.

Here, we tested the key prediction from the migratory divide hypothesis that hybrids have lower fitness than parental forms, focusing on survival as our fitness component. We used two datasets for this work – adults fitted with archival tags and juveniles fitted with radio tags. We used recapture and detection rates (respectively) as our proxy for survival. Using morphological and behavioural phenotypes from the juvenile dataset, we then asked whether variation in survival could be explained by selection against intermediate phenotypes or mismatch between phenotypes.

## Methods

### Estimating survival

Survival on migration was estimated separately for adults and juveniles. Adult birds show high site fidelity, returning to the same territory every year. Accordingly, we tracked this age class with archival tags (n = 285; light-level geolocators or GPS tags), fitting birds with tags during the breeding season (June 2010 – 2012 and 2019 – 2022, Table S1; Delmore and Irwin 2014) and surveying these and neighbouring territories the next year for returned birds. Birds that returned to the breeding grounds were counted as survived.

Juvenile birds disperse from their natal grounds, preventing recapture in the subsequent year. Accordingly, we relied on the Motus network (autonomous radio stations) to track this age class (n = 355), fitting birds with radio tags during fall migration, when they were fully fledged (August and September of 2019 – 2022, Table S1). Birds fitted with radio tags are detected when they fly within ∼20 km of a Motus station. We relied on existing Motus stations and established a fence of our own stations across the entire width of the hybrid zone in southern British Columbia (Fig. 1A). We released juveniles north of this fence each year; survival probabilities were modelled based on Motus station detections (see below for details).

We downloaded detection data from the Motus website (www.motus.org) on June 29, 2023 (Delmore and Easton 2019). We removed less reliable detections (detection run length < 3) first and inspected each remaining detection manually by plotting each bird’s movement during migration. This inspection was performed blind to ancestry or other information about each bird, and biologically improbable sightings were removed. We excluded data from five stations in northeastern North America with several faulty detections (LacEdouard-Champs, Lambs Gap, Lockoff, Tadoussac – Andr?, and Triton2). We also excluded all detections east of 70°W, detections below 20°N in June to August and above 35°N in December to February.

We constructed two capture histories for juveniles using Motus detection data – one for a two-state capture-recapture model and another for a multistate capture-recapture model. In both capture histories, each day was one time step, beginning with the day a bird was tagged. Capture histories were 300 days long to cover the data collection period for birds tagged in August 2022. For the two-state model capture history, if a bird was detected in a given calendar day, it was coded as “1” (vs. “0”). For the multistate model capture history, a bird detected above 40°N within 150 days of tagging was recorded as being in state one (pre-migratory), a bird detected below 40°N was recorded as being in state two (migratory), and a bird detected above 40°N over 150 days after tagging was considered to be in state three (post-migratory). The 150 day cutoff falls in late January, when birds are on non-breeding sites between the end of fall and start of spring migration (Delmore and Irwin 2014). We recorded birds detected after day 300 as detected (two-state model) or in state three (multistate model) on day 300.

### Tissue sample collection and genotype imputation

We collected tissue samples when tagging birds, using protocols approved by the Institutional Animal Care and Use Committee at Texas A&M (IACUC 2021-0068). All sampling was performed in accordance with relevant guidelines and regulations, including permits obtained from Environment and Climate Change Canada (10921), the Alaska Department of Fish and Game (20-1134, 21-117, 22-083), the Washington Department of Fish and Wildlife (20-104, 21-076, 22-061), the United States Department of the Interior (24199), the United States Fish and Wildlife Service (MB65923D), and the United States Department of Agriculture (USDA - 137701). DNA was extracted from blood using a standard phenol-chloroform protocol. We prepared libraries following a protocol modified from Picelli et al. (2014) and Schumer et al. (2018), then sequenced genomes to low coverage (average 3.8x). We aligned sequences to the reference genomes from each subspecies (assembly details in *Supplementary Methods*) using bwa and filtered to remove low quality sequences (Li and Durbin 2009). We used bcftools to call an initial set of SNPs (--min-BQ 20, --min-483 MQ 20, %QUAL>500, --skip-variants indels). Genotypes were then imputed using a hidden Markov model that estimates individuals’ haplotype probabilities, implemented through STITCH (see *Supplementary Methods*; Fig. S1; Davies et al. 2016).

### Ancestry and heterozygosity estimates

We filtered out SNPs with a minor allele frequency less than 5%, missing data greater than 25%, and more than two alleles. We applied linkage pruning with plink (50 10 0.1) and removed SNPs not in Hardy-Weinberg equilibrium (0.0001). We then removed SNPs on the Z and W chromosomes (scaffolds 7, 25, 26, 29) with vcftools (Danecek et al. 2011). Using the filtered dataset, we then estimated ancestry and heterozygosity in hybrid individuals. First, we needed to identify loci that distinguish the two subspecies. To do this, we estimated Weir-Cockerham F_ST_ using ten coastal and eight inland birds from our reference panel birds with vcftools (Danecek et al. 2011). We identified divergent sites as those with an F_ST_ greater than 0.94. Next, we estimated parental allele frequencies at each divergent site using the remaining reference panel birds (ten coastal and eight inland birds). Parental allele frequencies of 1 or 0 are likely to be the result of sampling error rather than true fixed differences between the subspecies, so parental allele frequencies above 0.98 and below 0.02 were adjusted to those values.

All analyses were performed in R version 4.2.3 and run separately for the adult and juvenile datasets (R Core Team 2022). We estimated ancestry and heterozygosity for each sample with the HIest R package, using allele counts estimated from the imputed genotypes (Fitzpatrick 2012). Ancestry is an estimate of the proportion of inland alleles, ranging from zero to one. Heterozygosity is the proportion of loci heterozygous for parental alleles. A coastal subspecies bird would have an ancestry of 0 and heterozygosity of 0, while an F_1_ hybrid would be expected to have an ancestry of 0.5 and a heterozygosity of 1 and an F_2_ hybrid would be expected to have an ancestry of 0.5 and a heterozygosity of 0.5 (Fig. 2B).

**Fig. 2.**
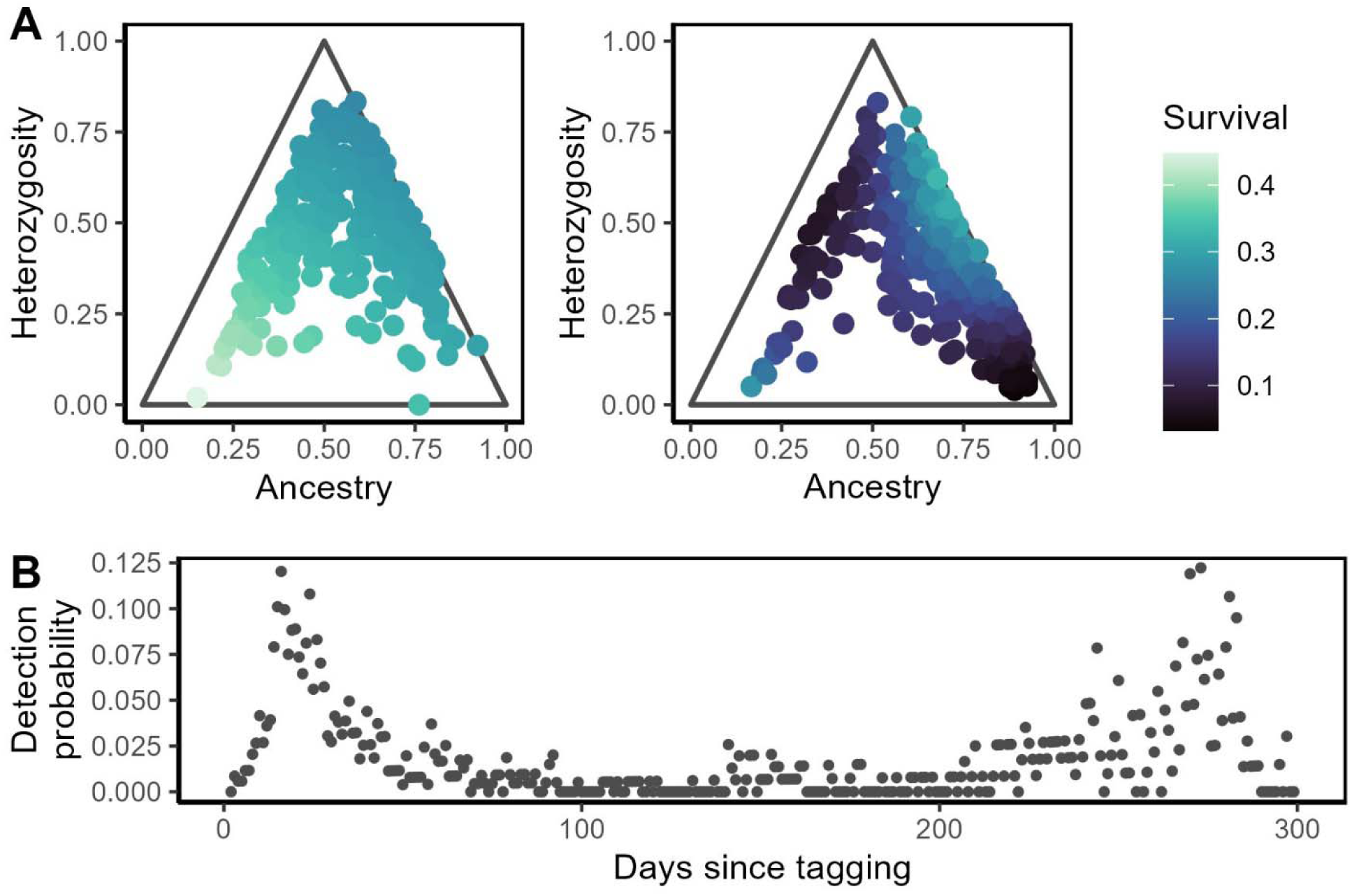
Survival by genomic class and detection across the annual cycle. (A) Relationship between survival, genetic ancestry, and heterozygosity for adults (left) and juveniles (right). Colour gradients represent fitted estimates of survival probabilities, from generalized linear mixed models (adults) or CJS models (juveniles). (B) Temporal variation in juvenile detection probability. Each point shows the probability of detection for a time point, based on CJS models and averaged over year, sex, ancestry, and heterozygosity.

### Survival by ancestry and heterozygosity

We tested the effects of genomic ancestry and heterozygosity on adult survival by fitting log-link generalized linear mixed models, with survival (recapture) as a binary response variable. Ancestry, heterozygosity, and their interaction were included in the model as fixed effects and release year and sex as random effects. Because migrating juvenile birds are detected across the Motus station network at multiple time points, we fit a Cormack-Jolly-Seber (CJS) model to estimate both survival and probability of detection based on the capture histories of juvenile birds (Bonner and Schwarz 2006; Lebreton et al. 2009). We fit a CJS model using the “hmmCJS” model from the marked R package, which implements a Bayesian Markov Chain Monte Carlo approach, with ancestry, heterozygosity, and their interaction as predictors of survival, and time step, release year, sex, ancestry, and heterozygosity as predictors of detection (Laake et al. 2013). We then fit a multistate CJS model to estimate survival, probability of transition between three migratory states (pre-migratory, migratory/overwintering, post-migratory), and probability of detection. The multistate CJS model was fit using the “hmmMSCJS” model, with ancestry, heterozygosity and their interaction as predictors of both survival and migratory state, and time step, release year, sex, ancestry, and heterozygosity as predictors of detection.

For juvenile birds, we then estimated the selection coefficient for hybrid relative to pure, parental subspecies individuals. We estimated survival as the average of predicted survival values among the observed ancestry and heterozygosity values corresponding to F_1_’s (heterozygosity > 0.75) and the subspecies (coastal: ancestry < 0.25, heterozygosity < 0.25; inland: ancestry > 0.75, heterozygosity < 0.75). We then estimated average survival for coastal backcrosses (ancestry < 0.4, 0.25 < heterozygosity < 0.75) and inland backcrosses (ancestry > 0.6, 0.25 < heterozygosity < 0.75). Selection coefficients were calculated as 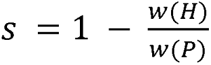 where *w(H)* was predicted hybrid survival (F_1_, coastal backcross, or inland backcross) and *w(P)* was the average of the predicted survival for the parental subspecies.

### Phenotypes

We tested for a relationship between phenotypes and survival in juveniles only, as migratory phenotype data could only be obtained for adult birds that were recaptured (i.e., survived). We focused on six migratory phenotypes. Four of these were morphological phenotypes: Kipp’s distance (distance between the first secondary feather and the longest primary), wing chord length, tail length, and tarsus length. Two were related to behaviour: migratory timing (date the bird crossed the southern BC transect on fall migration) and orientation (bearing on fall migration from the release site to a 30km radius). Morphological phenotypes were measured in the field and behavioural phenotypes were extracted from data recorded by Motus radio stations. To visualize how and whether phenotypes were correlated, we constructed phenotype correlation networks following Wilkins et al. (2015). We used Spearman’s ρ correlations, applied bootstrapping to generate 95% confidence intervals for correlations, and removed correlations with confidence intervals overlapping 0.

Because we observed high correlations between some phenotypes in the phenotype correlation network (Fig. 3), we first conducted a principal components analysis. To maximize the variance among axes, we applied a varimax rotation to the first three principal components (PCs; Fig. S2), which all had eigenvalues greater than 1 and cumulatively explained 71.4% of the variance. We tested the prediction that intermediate phenotypes would exhibit reduced survival relative using separate CJS models. Specifically, we evaluated whether survival depended on each of the first three PC axes, including both linear and quadratic terms, with time, release year, sex, ancestry, and heterozygosity as predictors of detection probability.

**Fig. 3.**
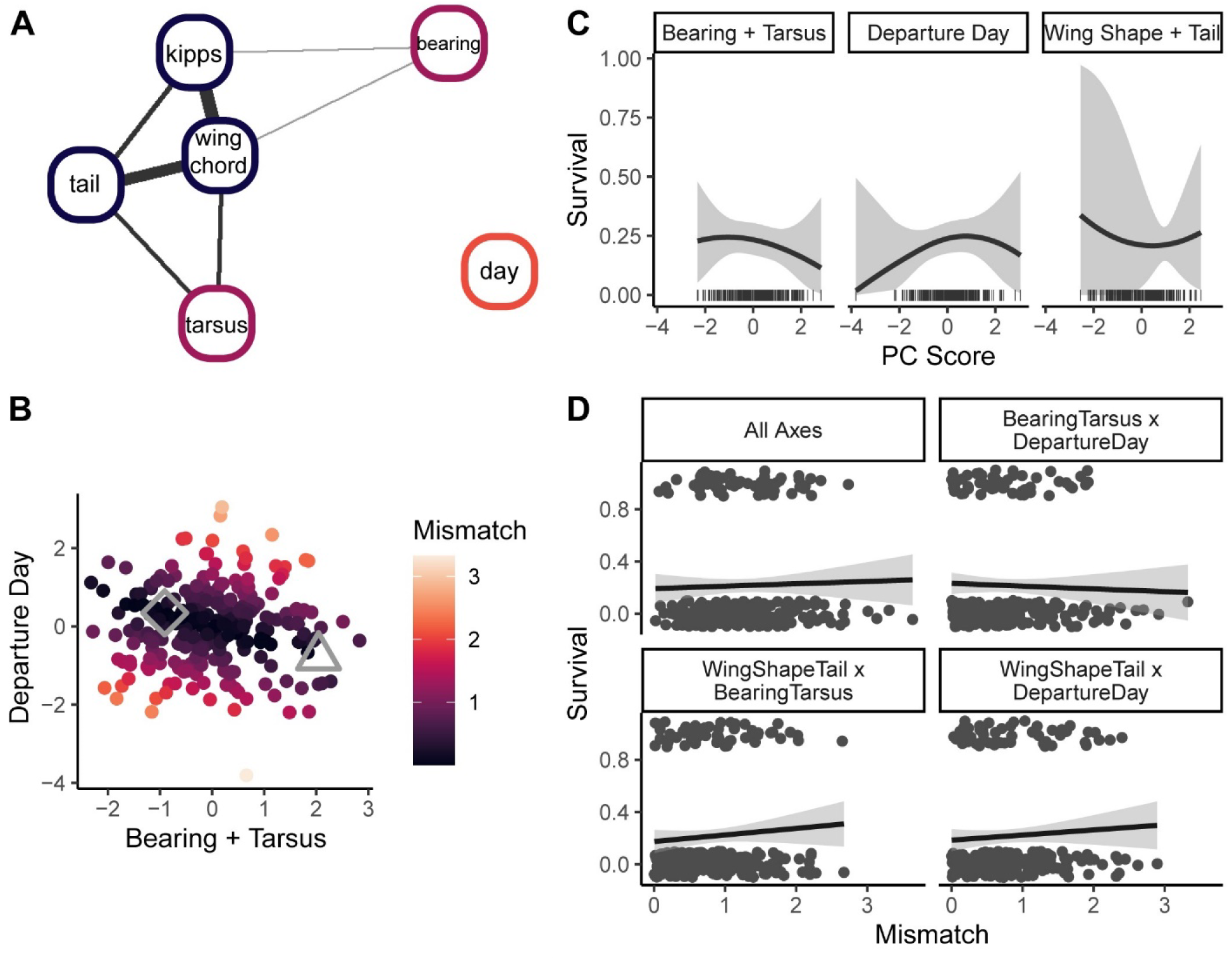
Relationship between survival and migratory phenotypes. (A) Network of phenotype correlations. Colours indicates PC axis along which each phenotype had the highest loading in a PCA of phenotypic variables among juvenile Swainson’s thrushes. PC1 primarily represented wing shape and tail length (green), PC2 represented fall migratory bearing and tarsus length (orange), and PC3 represented day of departure for fall migration (yellow). The width of edges indicates the strength of correlations. The minimum plotted correlation is ρ = 0.1. (B) Example phenotypic mismatch between two PC axes. Each point represents one individual. The open triangle and diamond (grey) indicate the estimated parental phenotype for the coastal and inland subspecies. (C & D) Relationships between survival on migration and (C) phenotypes and (D) phenotypic mismatch.. Lines represent model-based estimates of survival probabilities from CJS models and shaded regions represent confidence limits. No slopes (quadratic or linear) were significantly different from zero. Each tick along the x-axis represents the PC value for one bird.

For each individual, we estimated mismatch between each pair of PC axes as the Euclidean distance of the an individual from the line formed between the estimated parental values, following Chhina et al. (2022). We estimated parental values by fitting linear models with each PC axis as the response and ancestry as the predictor and extracting the estimated value where ancestry is 0 (coastal) or 1 (inland). To test the prediction that survival would decline with mismatch between phenotypes, we used separate CJS models with pairwise mismatch as the predictor of survival and time, release year, sex, ancestry, and heterozygosity as predictors of detection probability.

## Results

### Survival reduced in juvenile but not adult hybrids

We fitted 285 geolocator tags to adult birds during the breeding season and recaptured 89 in a subsequent year. There was minimal variation in survival with genomic class and no evidence for reduced survival (i.e., recapture) on migration in hybrid relative to parental adults (Fig. 2A). Using a generalized linear model testing for an effect of ancestry and heterozygosity on adult survival during migration, we found that survival varied minimally with ancestry (χ^2^_1_ = 0.59, p = 0.44), heterozygosity (χ^2^_1_ = 0.59, p = 0.44), and their interaction (χ^2^_1_ < 0.004, p = 0.94; Fig. 2A).

We fitted 355 radio tags to juveniles at the start of migration and 287 tags were detected by at least one station. Unlike with adults, we did find variation in survival (i.e., radio station detection) by genomic class (Fig. 2A). In the two state (survived and dead) model, survival declined with both ancestry (slope: -1.69 [confidence limit: -3.16, -0.23]), indicating reduced survival of more inland genotypes, and heterozygosity (-3.79 [-6.77, -0.82]), indicating lower survival of hybrids. There was additionally a significant interaction between ancestry and heterozygosity (7.67 [2.94, 12.41]), reflecting an asymmetrical relationship between survival and heterozygosity among birds with more coastal and more inland genomic backgrounds. Coastal backcrosses and first generation hybrids exhibited the lowest survival, along with the inland subspecies, while the coastal subspecies and inland backcrosses survived comparatively well (Fig. 2A). Detection probability varied among time points (Fig. 2B), peaking during fall and spring migration. Detection probability also varied among release years, increasing through time (year 2020: 0.91 [0.56, 1.25]; year 2021: 1.11 [0.77, 1.45]; year 2022: 1.50 [1.18, 1.82]), but not between sexes (-0.07 [-0.24, 0.10]). Birds with more inland ancestry had higher chances of detection (0.64 [0.15, 1.14]), but there was minimal detection probability variation with heterozygosity (0.28 [-0.18, 0.74]). We additionally fit a multistate model that estimated transitions between migratory stages (pre-migratory, migratory and overwintering, and post-migratory) in addition to survival and detection. Effects of ancestry, heterozygosity, and their interaction on transitions between migratory stages were in the same direction as their effects on survival in the two state model, but non-significant (ancestry: -1.33 [-3.36, 0.70]; heterozygosity: -1.83 [-6.32, 2.65]; ancestry x heterozygosity: 1.73 [-5.21, 8.67]).

To estimate the strength of selection against hybrid classes (i.e., F_1_’s, coastal and inland backcrosses), we estimated survival using fitted estimates from the two-state CJS model for corresponding ancestry and heterozygosity values (Fig. 2A). Juvenile F_1_ hybrids had a predicted probability of survival of 18%, while the parental subspecies had a predicted survival of 22%.

Predicted survival was 9% for coastal backcrosses and 24% for inland backcrosses. We obtained selection coefficients suggesting that there was selection against F_1_ hybrids and coastal backcrosses, but no evidence for selection against inland backcrosses. The selection coefficient for F_1_ hybrids relative to the parental subspecies was 0.15, meaning that an F_1_ hybrid had a chance of survival that was 85% that of the parental subspecies. The selection coefficient relative to the parental subspecies was 0.58 for coastal backcrosses. Contrary to predictions, inland backcrosses survived better on average than the parental subspecies, so the selection coefficient was -0.10 for inland backcrosses.

### No evidence for phenotypic selection related to migration

In juveniles, wing and tail phenotypes were tightly correlated, while behavioural phenotypes (fall bearing and fall day of departure) were only weakly correlated to morphological measures (Fig. 3A). There was also no correlation between fall day of departure and fall bearing. We summarised correlated phenotypic variation along four PC axes, each of which was renamed according to the loading of phenotypes along that axis (Fig. S2), with PC1 renamed Wing Shape & Tail, PC2 renamed Bearing & Tarsus, and PC3 renamed Departure Day. Contrary to the prediction of selection against intermediate phenotypes, there was no evidence for a linear or quadratic relationship between phenotype and survival along any of the four PC axes (Table 1; Fig. 3C).

**Table 1.** Effects of phenotypes on survival.

| **Predictor** |  | **Slope** | **Confidence Limit** |
| --- | --- | --- | --- |
| *Wing Shape + Tail* | *linear* | -0.04 | -0.89, 0.82 |
|  | *quadratic* | 0.04 | -0.13, 0.21 |
| *Bearing + Tarsus* | *linear* | -0.06 | -0.21, 0.10 |
|  | *quadratic* | -0.03 | -0.15, 0.09 |
| *Departure Day* | *linear* | 0.07 | -0.08, 0.23 |
|  | *quadratic* | -0.05 | -0.16, 0.06 |

We also tested for selection against phenotypic mismatch, which represents the distance between an individual’s phenotype and the slope between coastal and inland values for two phenotypes (Fig. 3B; Fig. S3). A hybrid with an intermediate value for both phenotypes would have low mismatch, but a hybrid with a coastal-like value for one phenotype and an inland-like value for the other would have high mismatch. Similar to results for intermediate phenotypes, we did not document selection against phenotypic mismatch, as survival did not decline significantly with increasing mismatch between any pairs of PC axes (Table 2; Fig. 3D; Fig. S3).

**Table 2.** Effects of phenotypic mismatch on survival.

| **Predictor** | **Slope** | **Confidence Limit** |
| --- | --- | --- |
| *Mismatch All* | 0.04 | -0.32, 0.39 |
| *Bearing + Tarsus × Departure Day* | -0.07 | -0.32, 0.19 |
| *Wing Shape + Tail × Bearing + Tarsus* | 0.12 | -0.16, 0.39 |
| *Wing Shape + Tail × Departure Day* | 0.08 | -0.18, 0.33 |

## Discussion

Migratory divides are predicted to be an important mechanism underlying ecological speciation, serving as an extrinsic postzygotic barrier to gene flow. The Swainson’s thrush is one of the only systems where direct tracking data have shown hybrids take intermediate routes on migration, supporting the first prediction of the migratory divide hypothesis (Delmore and Irwin 2014). Here, we tested whether hybrids experienced reduced survival on migration relative to parental forms using return/detection rates as a proxy for this fitness component. We then attempted to link reductions in survival to either intermediate or mismatched migratory phenotypes. As expected, juvenile hybrids, especially F_1_’s and coastal backcrosses, survived at lower rates than pure, parental individuals. In contrast, there was no evidence for reduced survival in hybrids at the adult life stage. Adult birds were fitted with archival tags after completing migration successfully at least once, so variation in survival across genomic backgrounds might not be expected at this life stage. Although some form of selection reduced the fitness of juvenile hybrids once they left the breeding grounds, this pattern was not explained by reduced survival of intermediate migratory phenotypes or mismatched phenotypes.

Ecological barriers are likely critical for maintaining divergence between many species, but extrinsic selection against hybrids has been demonstrated in only a few systems, such as threespine stickleback (Arnegard et al. 2014; Thompson et al. 2022), crossbills (Benkman 2003), and *Heliconius* butterflies (Merrill et al. 2012). Migratory divides are common in nature (Irwin and Irwin 2005; Møller et al. 2011; Rohwer and Irwin 2011), and our results indicate that selection against hybrids on migration likely contributes to speciation between Swainson’s thrush subspecies. Previous estimates from cline theory indicated that moderate to strong selection against hybrids is maintaining the hybrid zone between the coastal and inland subspecies (Delmore and Irwin 2014). We obtained moderate selection coefficients for selection against F_1_ hybrids and coastal backcrosses, suggesting reduced survival on migration contributes substantially to selection against hybrids and maintenance of the narrow hybrid zone (Barton and Hewitt 1985; Irwin 2020). If selection were intrinsic rather than linked to migration, specifically, then hybrids would be expected to show similarly reduced survival in a lab environment and during other parts of the annual cycle. Instead, wild-caught hybrid juveniles raised in captivity over the fall and spring migration periods do not show reduced survival (Louder et al. 2024) and radio-tagged hybrids that survived migration were detected again after the breeding season (Fig. S5), suggesting that reduced hybrid survival is specific to migration.

In contrast to the initial formulation of the migratory divide hypothesis, which emphasized that intermediate ancestry will exhibit low fitness relative to parental forms (Helbig 1991; Irwin and Irwin 2005), we found asymmetric selection across genomic ancestries. Specifically, coastal backcrosses and F_1_’s survived at low rates on migration while inland backcrosses survived well, relative to parental forms. Migration may represent an asymmetrical barrier to introgression in this system, in which coastal alleles should be able to persist in a mostly inland genomic background but inland alleles cannot do so in a mostly coastal genomic background. This asymmetry may have an ecological origin. Specifically, the inland subspecies has a broad range (Fig. 1) with suitable habitat and high connectivity across much of eastern North American (Justen et al. 2021). Introgression of coastal alleles into the inland subspecies may be beneficial if these birds take a shorter, more western route that is eastward enough to avoid the regions with lowest habitat connectivity. In contrast, the coastal subspecies only has a narrow region of suitable habitat (Fig 1.; Justen et al. 2021), meaning that even a relatively minor shift eastward may be enough for them to encounter obstacles and unsuitable stopover sites. Therefore, genomic background, migratory phenotype, and geography may interact to determine success on migration.

Migrating juvenile birds with more inland ancestry tended to have a higher probability of radio station detection than coastal birds. This likely reflects differences in the density of Motus receiver stations between the subspecies’ migration routes, given the high concentration of stations in eastern North America (Fig. 1). Similarly, the increase in detection probability across study years likely results from an increase through time in the number of active Motus stations, particularly in Central America and western North America (Taylor et al. 2017). However, this variability in detection probability likely does not account for the reduced survival inferred for migrating hybrid juveniles. If survival patterns were produced by ancestry-related variation in detection rates, we would expect to see lower survival in coastal birds than in coastal backcrosses and F1 hybrids and higher survival in inland birds than inland backcrosses. Instead, we see the opposite patterns, suggesting that genomic background does impact survival on migration.

While we have shown that some form of selection acted against hybrids while they were away from the breeding grounds, reduced fitness of some hybrid genotypes was not explained by selection against either intermediate phenotypes or phenotypic mismatch. There are multiple possible explanations for the absence of this pattern. Reduced hybrid survival may not be a result of the same mechanism across all individuals. If some individuals experience a mismatch between their timing and destination of fall migration, while others take an intermediate route, and others have the wrong wing shape for their route, then all would have reduced survival but a trend across the population would not emerge. Additionally, it is possible that we are measuring the wrong phenotypes or using imperfect assessments of them. For example, several behavioural phenotypes that differ between the subspecies cannot be inferred with the radio tag dataset, such as total migration distance and duration and location of stopover and nonbreeding sites (Delmore and Irwin 2014). Other migratory phenotypes that may have contributed to reduced survival are infeasible to assay in the field, such as physiological or cellular processes (McWilliams et al. 2004). For example, captive Swainson’s thrush hybrids exhibit high levels of transgressive gene expression during migration seasons, which could contribute to reduced survived (Louder et al. 2024).

We found that selection on genomic ancestry during migration occurs primarily during the first year of migration and acts against some hybrid classes. However, open questions remain in this system, including which specific phenotypes or phenotypic interactions are under selection. Nonetheless, our results provide support for the idea that reduced survival of hybrids may contribute to speciation in a migratory divide. Migratory divides are common in nature and have been hypothesized to contribute to several cases of incipient speciation (Irwin and Irwin 2005; Irwin 2009; Delmore et al. 2015). Swainson’s thrushes are likely not the only system where hybrids experience reduced survival on migration. Some known migratory divides circumvent geographical features that would likely impose greater mortality on hybrids with intermediate routes than those encountered by Swainson’s thrushes. For example, divides between barn swallows and greenish warblers that breed in China involve navigation around the Tibetan Plateau and willow warblers that breed in Scandinavia take routes either east or west of the Mediterranean Sea and Sahara Desert (Irwin and Irwin 2005; Turbek et al. 2022; Sokolovskis et al. 2023). Therefore, migratory divides may constitute a relatively common and important form of extrinsic, post-zygotic isolation in migrating species.

## Acknowledgements

Funding for this research came from an NSF CAREER grant to KED (IOS-2143004), start-up funds from Texas A&M University, and an operating grant to WEE from CWS-ECCC. We greatly appreciate assistance in the field from members of the Delmore lab (especially Miranda Anderson, Hayley Madden, Scarlet Byron and Catherine Paul and several undergraduates), Todd Alleger, Zoe Crysler, John Klicka, and Kevin Epperly. We also thank landowners that facilitate our fence of Motus stations (project #280) in British Columbia and all collaborators contributing to the Motus Wildlife Tracking system. We thank Darren Irwin for comments on an earlier version of the manuscript.

## References

Albert, V., B. Jonsson, and L. Bernatchez. 2006. Natural hybrids in Atlantic eels (Anguilla anguilla, A . rostrata): evidence for successful reproduction and fluctuating abundance in space and time. Molecular Ecology 15:1903–1916.

Arnegard, M. E., M. D. McGee, B. Matthews, K. B. Marchinko, G. L. Conte, S. Kabir, N. Bedford, et al. 2014. Genetics of ecological divergence during speciation. Nature 511:307–311.

Barton, N. H., and G. M. Hewitt. 1985. Analysis of hybrid zones. Annual review of ecology and systematics. Vol. 16 16:113–148.

Benkman, C. W. 2003. Divergent selection drives the adaptive radiation of crossbills. Evolution 57:1176–1181.

Bonner, S. J., and C. J. Schwarz. 2006. An extension of the Cormack-Jolly-Seber model for continuous covariates with application to Microtus pennsylvanicus. Biometrics 62:142–149.

Bridge, E. S., K. Thorup, M. S. Bowlin, P. B. Chilson, R. H. Diehl, R. W. Fléron, P. Hartl, et al. 2011. Technology on the move: Recent and forthcoming innovations for tracking migratory birds. BioScience 61:689–698.

Chhina, A. K., K. A. Thompson, and D. Schluter. 2022. Adaptive divergence and the evolution of hybrid trait mismatch in threespine stickleback. Evolution Letters 6:34–45.

Danecek, P., A. Auton, G. Abecasis, C. A. Albers, E. Banks, M. A. DePristo, R. E. Handsaker, et al. 2011. The variant call format and VCFtools. Bioinformatics 27:2156–2158.

Davies, R. W., J. Flint, S. Myers, and R. Mott. 2016. Rapid genotype imputation from sequence without reference panels. Nature Genetics 48:965–969.

Delmore, K. E., and W. Easton. 2019. BC Interior Thrushes (Project280). Motus Wildlife Tracking System, Birds Canada.

Delmore, K. E., and D. E. Irwin. 2014. Hybrid songbirds employ intermediate routes in a migratory divide. Ecology Letters 17:1211–1218.

Delmore, K. E., H. L. Kenyon, R. R. Germain, and D. E. Irwin. 2015. Phenotypic divergence during speciation is inversely associated with differences in seasonal migration. Proceedings of the Royal Society B: Biological Sciences 282.

Dettman, J. R., C. Sirjusingh, L. M. Kohn, and J. B. Anderson. 2007. Incipient speciation by divergent adaptation and antagonistic epistasis in yeast. Nature 447:585–588.

Dieckmann, U., and M. Doebeli. 1999. On the origin of species by sympatric speciation. Nature 400:354–357.

Dingle, H. 2006. Animal migration: is there a common migratory syndrome? Journal of Ornithology 147:212–220.

Fitzpatrick, B. M. 2012. Estimating ancestry and heterozygosity of hybrids using molecular markers. BMC evolutionary biology 12:1–14.

Helbig, A. J. 1991. SELJ and SWLJmigrating Blackcap (Sylvia atricapilla) populations in Central Europe: Orientation of birds in the contact zone. Journal of Evolutionary Biology 4:657–670.

Irwin, D. E. 2009. Speciation: New Migratory Direction Provides Route toward Divergence. Current Biology 19:R1111–R1113.

Irwin, D. E. 2020. Assortative mating in hybrid zones is remarkably ineffective in promoting speciation. American Naturalist 195:E150–E167.

Irwin, D. E., and J. H. Irwin. 2005. Siberian migratory divides: the role of seasonal migration in speciation. Pages 27–40 in R. Greenberg and P. P. Marra, eds. Birds of Two Worlds: The Ecology and Evolution of Migration. Johns Hopkins University Press.

Justen, H., and K. E. Delmore. 2022. The genetics of bird migration. Current Biology 32:R1144–R1149.

Justen, H., J. A. Lee-Yaw, and K. E. Delmore. 2021. Reduced habitat suitability and landscape connectivity in a songbird migratory divide. Global Ecology and Biogeography 30:2043–2056.

Laake, J. L., D. S. Johnson, and P. B. Conn. 2013. marked: An R package for maximum likelihood and Markov Chain Monte Carlo analysis of capture-recapture data. Methods in Ecology and Evolution 4:885–890.

Lebreton, J. D., J. D. Nichols, R. J. Barker, R. Pradel, and J. A. Spendelow. 2009. Modeling Individual Animal Histories with Multistate Capture – Recapture Models. Advances in Ecological Research (1st ed., Vol. 41). Elsevier Ltd.

Li, H., and R. Durbin. 2009. Fast and accurate short read alignment with Burrows – Wheeler transform. Bioinformatics 25:1754–1760.

Louder, M. I. M., H. Justen, A. A. Kimmitt, K. S. Lawley, L. M. Turner, J. D. Dickman, and K. E. Delmore. 2024. Gene regulation and speciation in a migratory divide between songbirds. Nature Communications 15:1–14.

McWilliams, S. R., C. Guglielmo, B. Pierce, and M. Klaassen. 2004. Flying, fasting, and feeding in birds during migration: A nutritional and physiological ecology perspective. Journal of Avian Biology 35:377–393.

Merrill, R. M., R. W. R. Wallbank, V. Bull, P. C. A. Salazar, J. Mallet, M. Stevens, and C. D. Jiggins. 2012. Disruptive ecological selection on a mating cue. Proceedings of the Royal Society B: Biological Sciences 279:4907–4913.

Møller, A. P., J. M. Peralta-Sanchez, L. Z. Garamszegi, and J. J. Soler. 2011. Migratory divides and their consequences for dispersal, population size and parasite – host interactions. Journal of Evolutionary Biology 24:1744–1755.

Newton, I. 2006. Can conditions experienced during migration limit the population levels of birds? Journal of Ornithology 147:146–166.

Paradis, E., S. R. Baillie, W. J. Sutherland, and R. D. Gregory. 1998. Patterns of natal and breeding dispersal in birds. Journal of Animal Ecology 67:518–536.

Picelli, S., A. K. Bjorklund, B. Reinius, S. Sargasser, G. Winberg, and R. Sandberg. 2014. Tn5 transposase and tagmentation procedures for massively scaled sequencing projects. Genome Research 24:2033–2040.

R Core Team. 2022. R: A Language and Environment for Statistical Computing. R Foundation for Statistical Computing, Vienna, Austria.

Reppert, S. M., R. J. Gegear, and C. Merlin. 2010. Navigational mechanisms of migrating monarch butterflies. Trends in Neurosciences 33:399–406.

Robinson, R. A., C. M. Meier, W. Witvliet, M. Kéry, and M. Schaub. 2020. Survival varies seasonally in a migratory bird: Linkages between breeding and non-breeding periods. Journal of Animal Ecology 89:2111–2121.

Rohwer, S., and D. E. Irwin. 2011. Molt, orientation, and avian speciation. Auk.

Ruegg, K. 2008. Genetic, morphological, and ecological characterization of a hybrid zone that spans a migratory divide. Evolution 62:452–466.

Rundle, H. D., and P. Nosil. 2005. Ecological speciation. Ecology Letters 8:336–352.

Schluter, D. 2000. The Ecology of Adaptive Radiation. OUP, Oxford.

Schumer, M., C. Xu, D. L. Powell, A. Durvasula, L. Skov, C. Holland, J. C. Blazier, et al. 2018. Natural selection interacts with recombination to shape the evolution of hybrid genomes. Science 3684:eaar3684.

Scordato, E. S. C., C. C. R. Smith, G. A. Semenov, Y. Liu, M. R. Wilkins, W. Liang, A. Rubtsov, et al. 2020. Migratory divides coincide with reproductive barriers across replicated avian hybrid zones above the Tibetan Plateau. Ecology Letters 23:231–241.

Sergio, F., G. Tavecchia, A. Tanferna, J. Blas, G. Blanco, and F. Hiraldo. 2019. When and where mortality occurs throughout the annual cycle changes with age in a migratory bird: individual vs population implications. Scientific Reports 9:1–8.

Sokolovskis, K., M. Lundberg, S. Åkesson, M. Willemoes, T. Zhao, V. Caballero-lopez, and S. Bensch. 2023. Migration direction in a songbird explained by two loci. Nature Communications 14:165.

Taylor, P. D., T. L. Crewe, S. A. Mackenzie, D. Lepage, Y. Aubry, Z. Crysler, G. Finney, et al. 2017. The Motus Wildlife Tracking System: a collaborative research network. Avian Conservation & Ecology 12:8.

Thompson, K. A., C. L. Peichel, D. J. Rennison, M. D. McGee, A. Y. K. Albert, T. H. Vines, A. K. Greenwood, et al. 2022. Analysis of ancestry heterozygosity suggests that hybrid incompatibilities in threespine stickleback are environment dependent. PLoS Biology 20:1–19.

Turbek, S. P., D. R. Schield, E. S. C. Scordato, A. Contina, X. Da, Y. Liu, Y. Liu, et al. 2022. A migratory divide spanning two continents is associated with genomic and ecological divergence. Evolution 76:722–736.

Vali, U., P. Mirski, U. Sellis, M. Dagys, and G. Maciorowski. 2018. Genetic determination of migration strategies in large soaring birds: evidence from hybrid eagles. Proceedings of the Royal Society B: Biological Sciences 285:20180855.

Wilkins, M. R., D. Shizuka, M. B. Joseph, J. K. Hubbard, and R. J. Safran. 2015. Multimodal signalling in the North American barn swallow: A phenotype network approach. Proceedings of the Royal Society B: Biological Sciences 282.

