## Supplemental Methods and Figures for "Reduced hybrid survival in a migratory divide between songbirds"

**Supplementary Methods**

*Reference genome assembly*

The inland subspecies’ genome was assembled and annotated as part of the Vertebrate 10K genome project using PacBio, Hi-C, Bionano and Illumina short-read technologies and tissue from a female thrush captured in Kamloops, British Columbia (VGP assembly pipeline v. 1.5, https://www-ncbi-nlm-nih-gov.srv-proxy2.library.tamu.edu/assembly/GCF_009819885.2). We assembled the coastal subspecies genome using PacBio, Dovetail Chicago and Hi-C technologies and two female thrushes. DNA was extracted from both samples with Qiagen’s Genomic DNA Kit. We constructed PacBio and Chicago libraries using blood from one female captured in Vancouver, British Columbia. We constructed Hi-C libraries using muscle from the second female, captured in Haida Gwaii, British Columbia. PacBio libraries were sequenced on seven SMRT cells while Chicago and Hi-C libraries were sequenced together on a single lane of Illumina’s HiSeqX (2x150 bp). We assembled the PacBio data using canu and default settings then polished the assembly using two rounds of read mapping with Pilon (Walker et al. 2014; Koren et al. 2017). To scaffold the PacBio assembly, Dovetail Genomics used their Hi Rise software pipeline (Dovetail Genomics, LLC) with Chicago and Hi-C data. We then filled gaps using the PacBio data and PBJelly (English et al. 2012). The final assembly was 1.1 billion base pairs in length, comprising 221 scaffolds (N50 = 73.3 Mb) and 820 914 contigs (N50 = 40.8 Mb).

*Genotype imputation*

An initial set of SNPs for STITCH was created by filtering with bcftools (--min-BQ 20, --min-MQ 20, QUAL>500, --skip-variants indels; Li 2011; Danecek et al. 2021). STITCH was run in blocks of 1 Mb with a 100 kb buffer (Davies et al. 2016). The program was initiated using the pseudoHaploid model, with values of 80 for ancestral haplotypes (K) and 500 for number of generations since the population was founded (nGen), then switched over to the diploid model after 36 EM. We assessed the accuracy of imputation using a cross-validation approach, randomly selecting four hybrids that were originally sequenced to low-coverage to be sequenced to high coverage. We used samtools to randomly subsample reads from each individual to mimic low-coverage sequencing (at 1, 2, 4 and 7x coverage) and compared genotypes called using high coverage data (i.e., “true” genotypes) to those estimated using low-coverage data. Comparisons were made using the squared linear correlation coefficient at a minor allele cutoff of 0.5, with R^2^ values averaged across sites and samples to obtain a single summary for each coverage. Figure S1 shows these results, with accuracy starting at ~0.95 at 1x coverage and increasing to ~0.975 at 4x. To identify divergent SNPs between the two subspecies, a reference panel of fourteen birds of each subspecies were sequenced to high coverage (average 18x), using whole genome resequencing libraries prepared with Nextera (Illumina) DNA Flex 744 Library Prep kits.

**References**

Danecek, P., J. K. Bonfield, J. Liddle, J. Marshall, V. Ohan, M. O. Pollard, A. Whitwham, et al. 2021. Twelve years of SAMtools and BCFtools. GigaScience 10:1–4.

Davies, R. W., J. Flint, S. Myers, and R. Mott. 2016. Rapid genotype imputation from sequence without reference panels. Nature Genetics 48:965–969.

English, A. C., S. Richards, Y. Han, M. Wang, V. Vee, J. Qu, X. Qin, et al. 2012. Mind the Gap: Upgrading Genomes with Pacific Biosciences RS Long-Read Sequencing Technology. PLoS ONE 7:e47768.

Koren, S., B. P. Walenz, K. Berlin, J. R. Miller, N. H. Bergman, and A. M. Phillippy. 2017. Canu: Scalable and accurate long-read assembly via adaptive κ-mer weighting and repeat separation. Genome Research 27:722–736.

Li, H. 2011. A statistical framework for SNP calling, mutation discovery, association mapping and population genetical parameter estimation from sequencing data. Bioinformatics 27:2987–2993.

Walker, B. J., T. Abeel, T. Shea, M. Priest, A. Abouelliel, S. Sakthikumar, C. A. Cuomo, et al. 2014. Pilon: An integrated tool for comprehensive microbial variant detection and genome assembly improvement. PLoS ONE 9:e112963.

**Supplementary Figures**

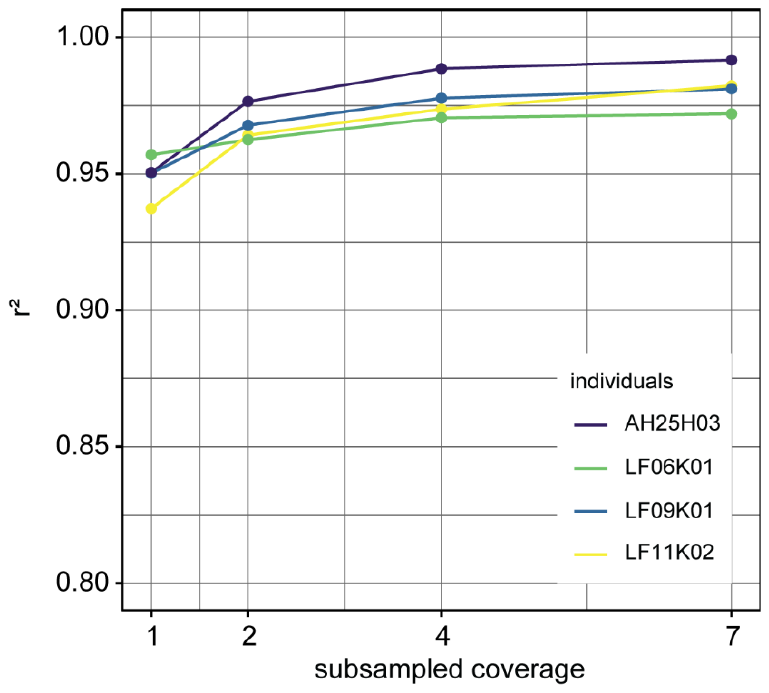

Fig. S1. Showing the accuracy of genotype imputation by STITCH. Squared correlation coefficients between imputed and actual genotypes for four hybrids sequenced to high coverage.

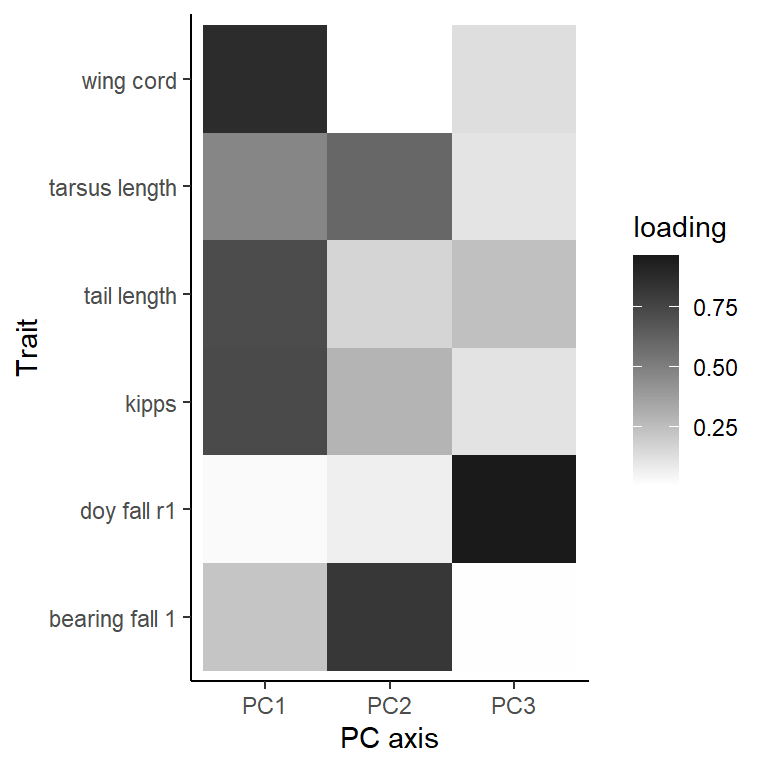

Fig. S2. PC loadings for each phenotype. PC axes were named according to the phenotypes with the highest loadings along that axis, with PC1 renamed Wing / Tail Length, PC2 renamed Bearing / Tarsus, PC3 renamed Wing Shape, and PC4 renamed Departure Day.

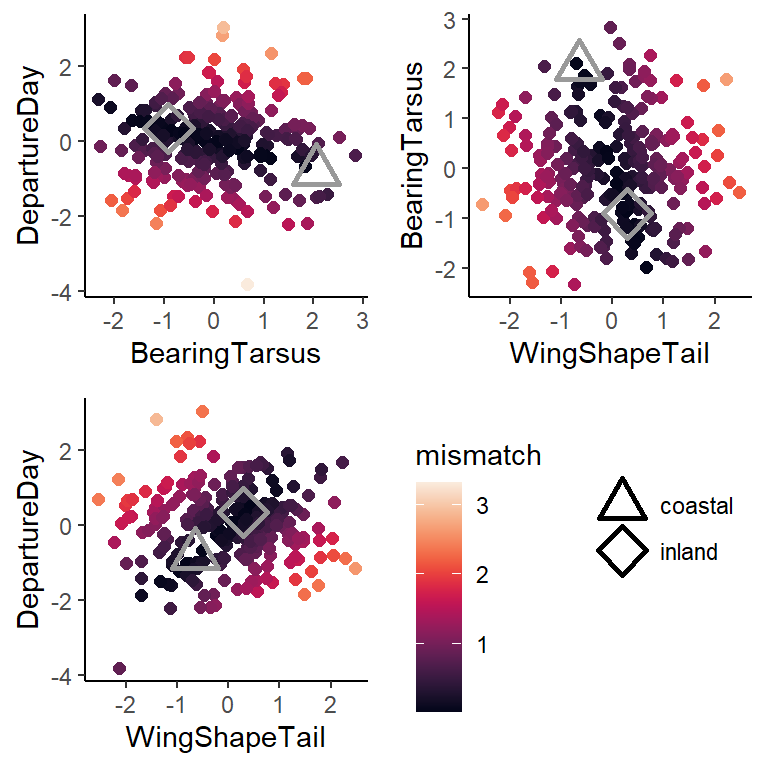

Fig. S3. Phenotypic mismatch in juveniles. Each point represents one individual. The open triangle and diamond indicate the estimated parental phenotype for the coastal and inland subspecies.

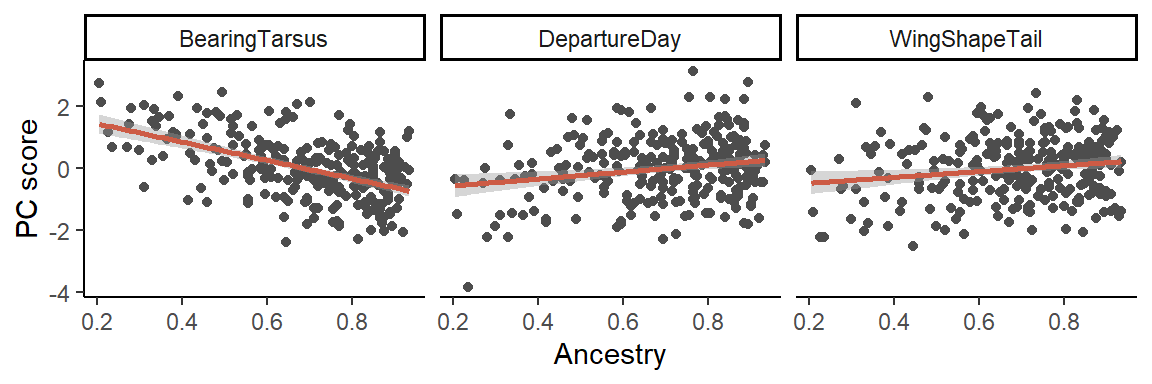

Fig. S4. Relationships between ancestry and PC scores. Each point represents one bird. Shaded regions represent 95% confidence intervals.

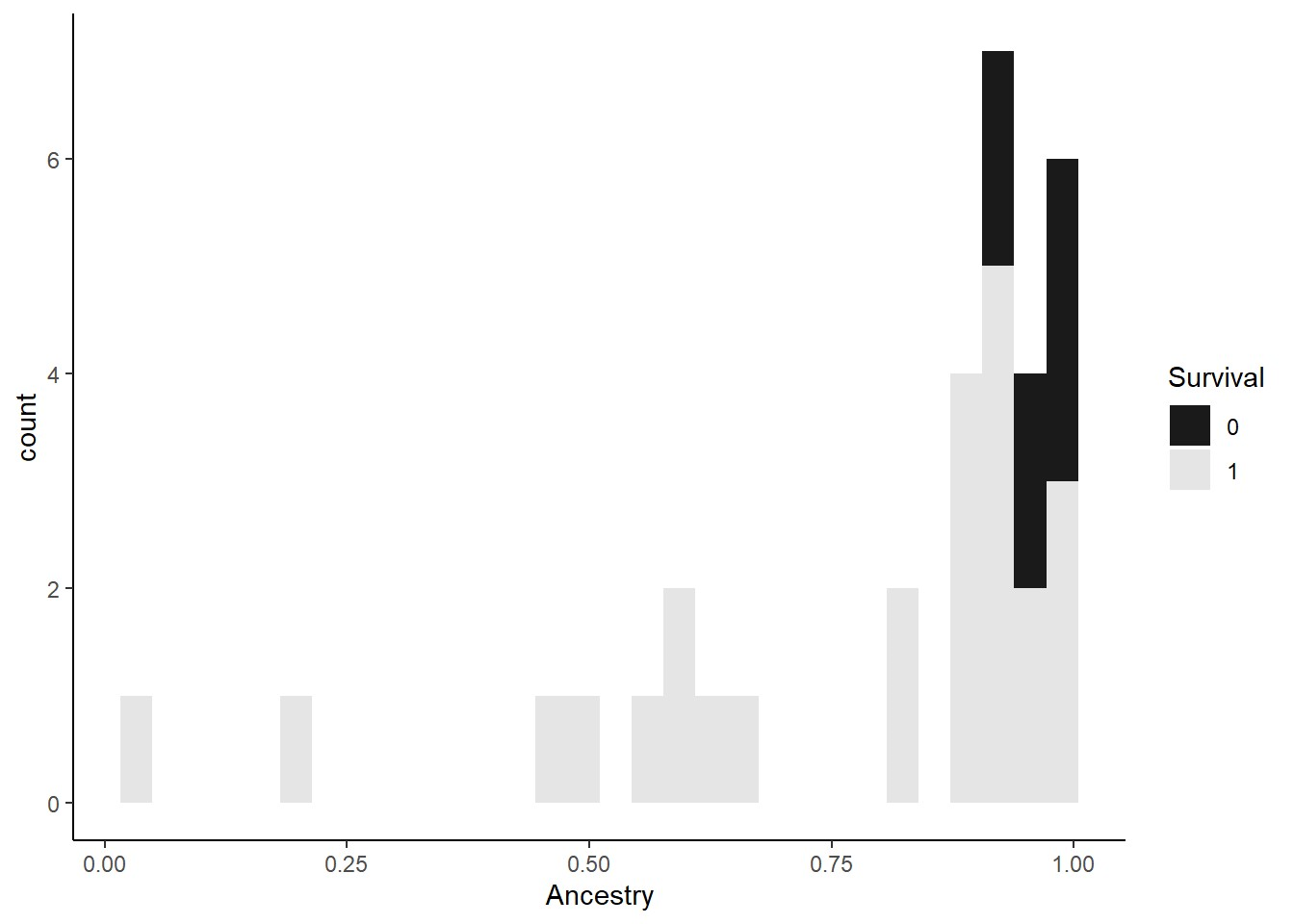

Fig. S5. Survival of juvenile hybrids over the breeding season. Juvenile birds released in 2021 or earlier that were detected in the breeding grounds (latitude > 40°N) in the spring are plotted, with “1” indicating birds that were detected again after the breeding season (August or later) and “0” indicating birds that were not detected again.

**Supplementary Tables**

Table S1. Tagging and sequence availability information for each bird including in the study.

| reference | release site | GPS coordinates | tag type | SRA BioProject |
| --- | --- | --- | --- | --- |
| AF12H01 | Hope | 49.38 N 121.32 W | archival | PRJNA1024534 |
| AF12H02 | Hope | 49.38 N 121.32 W | archival | PRJNA979932 |
| AF14H01 | Pemberton | 50.22 N 122.89 W | archival | PRJNA1024534 |
| AF14H04 | Pemberton | 50.37 N 122.87 W | archival | PRJNA1024534 |
| AF14H06 | Pemberton | 50.37 N 122.86 W | archival | PRJNA1024534 |
| AF16H01 | Pemberton | 50.22 N 122.89 W | archival | PRJNA1024534 |
| AF16H02 | Pemberton | 50.22 N 122.89 W | archival | PRJNA1024534 |
| AF16H03 | Pemberton | 50.22 N 122.89 W | archival | PRJNA979932 |
| AF16H04 | Pemberton | 50.37 N 122.86 W | archival | PRJNA979932 |
| AF17H01 | Pemberton | 50.37 N 122.86 W | archival | PRJNA979932 |
| AF17H02 | Pemberton | 50.37 N 122.86 W | archival | PRJNA1024534 |
| AF17H03 | Pemberton | 50.37 N 122.86 W | archival | PRJNA1024534 |
| AF17H04 | Pemberton | 50.22 N 122.89 W | archival | PRJNA979932 |
| AF19H01 | Pemberton | 50.37 N 122.86 W | archival | PRJNA1024534 |
| AF20H01 | Hope | 49.47 N 121.25 W | archival | PRJNA1024534 |
| AF20H02 | Hope | 49.47 N 121.25 W | archival | PRJNA979932 |
| AF20H03 | Hope | 49.47 N 121.25 W | archival | PRJNA1024534 |
| AF20H04 | Hope | 49.47 N 121.25 W | archival | PRJNA979932 |
| AF21H01 | Hope | 49.47 N 121.25 W | archival | PRJNA979932 |
| AF21H02 | Hope | 49.47 N 121.25 W | archival | PRJNA979932 |
| AF21H03 | Hope | 49.46 N 121.25 W | archival | PRJNA979932 |
| AF21H04 | Hope | 49.39 N 121.32 W | archival | PRJNA1024534 |
| AF22H01 | Hope | 49.48 N 121.25 W | archival | PRJNA979932 |
| AF22H02 | Hope | 49.48 N 121.25 W | archival | PRJNA1024534 |
| AF22H03 | Hope | 49.48 N 121.25 W | archival | PRJNA1024534 |
| AF22H04 | Hope | 49.48 N 121.25 W | archival | PRJNA1024534 |
| AF22H05 | Hope | 49.49 N 121.25 W | archival | PRJNA1024534 |
| AF22H06 | Hope | 49.48 N 121.25 W | archival | PRJNA1024534 |
| AF22H07 | Hope | 49.38 N 121.32 W | archival | PRJNA979932 |
| AF23H01 | Hope | 49.48 N 121.25 W | archival | PRJNA1024534 |
| AF23H02 | Hope | 49.47 N 121.25 W | archival | PRJNA979932 |
| AF23H03 | Hope | 49.48 N 121.25 W | archival | PRJNA1024534 |
| AF23H04 | Hope | 49.46 N 121.25 W | archival | PRJNA1024534 |
| AF23H05 | Hope | 49.48 N 121.25 W | archival | PRJNA1024534 |
| AF24H01 | Hope | 49.49 N 121.24 W | archival | PRJNA1024534 |
| AF24H02 | Hope | 49.49 N 121.24 W | archival | PRJNA1024534 |
| AF24H03 | Hope | 49.38 N 121.32 W | archival | PRJNA1024534 |
| AF26H01 | Hope | 49.48 N 121.25 W | archival | PRJNA1024534 |
| AF26H02 | Hope | 49.48 N 121.24 W | archival | PRJNA1024534 |
| AF26H03 | Hope | 49.47 N 121.25 W | archival | PRJNA1024534 |
| AF26H04 | Hope | 49.48 N 121.24 W | archival | PRJNA1024534 |
| AF26H05 | Hope | 49.47 N 121.25 W | archival | PRJNA1024534 |
| AF27H01 | Pemberton | 50.34 N 122.75 W | archival | PRJNA1024534 |
| AF27H02 | Pemberton | 50.34 N 122.75 W | archival | PRJNA979932 |
| AF27H03 | Pemberton | 50.34 N 122.74 W | archival | PRJNA1024534 |
| AF27H04 | Pemberton | 50.34 N 122.75 W | archival | PRJNA1024534 |
| AF28H01 | Pemberton | 50.33 N 122.75 W | archival | PRJNA1024534 |
| AF28H02 | Pemberton | 50.34 N 122.74 W | archival | PRJNA979932 |
| AF28H03 | Pemberton | 50.34 N 122.74 W | archival | PRJNA979932 |
| AF29H01 | Pemberton | 50.34 N 122.75 W | archival | PRJNA1024534 |
| AF29H02 | Pemberton | 50.34 N 122.75 W | archival | PRJNA979932 |
| AF29H03 | Pemberton | 50.34 N 122.75 W | archival | PRJNA979932 |
| AF30H01 | Pemberton | 50.34 N 122.74 W | archival | PRJNA1024534 |
| AF30H03 | Pemberton | 50.34 N 122.74 W | archival | PRJNA1024534 |
| AG01H01 | Pemberton | 50.33 N 122.74 W | archival | PRJNA1024534 |
| AG01H02 | Pemberton | 50.33 N 122.74 W | archival | PRJNA1024534 |
| AG01H03 | Pemberton | 50.34 N 122.74 W | archival | PRJNA1024534 |
| AG01H04 | Pemberton | 50.33 N 122.74 W | archival | PRJNA1024534 |
| AH17K18 | Pemberton | 50.35 N 122.83 W | radio | PRJNA979932 |
| AH18K02 | Pemberton | 50.35 N 122.83 W | radio | PRJNA979932 |
| AH22K02 | Pemberton | 50.35 N 122.83 W | radio | PRJNA979932 |
| AH22K04_S154_L001 | Pemberton | 50.35 N 122.83 W | radio | PRJNA1024534 |
| AH22K09 | Pemberton | 50.35 N 122.83 W | radio | PRJNA979932 |
| AH22K11_S155_L001 | Pemberton | 50.35 N 122.83 W | radio | PRJNA1024534 |
| AH22K13 | Pemberton | 50.35 N 122.83 W | radio | PRJNA979932 |
| AH22K14_S156_L001 | Pemberton | 50.35 N 122.83 W | radio | PRJNA1024534 |
| AH22K15 | Pemberton | 50.35 N 122.83 W | radio | PRJNA979932 |
| AH24H02 | Pemberton | 50.35 N 122.83 W | radio | PRJNA979932 |
| AH24H03 | Pemberton | 50.35 N 122.83 W | radio | PRJNA979932 |
| AH24H04 | Pemberton | 50.35 N 122.83 W | radio | PRJNA979932 |
| AH24H05 | Pemberton | 50.35 N 122.83 W | radio | PRJNA979932 |
| AH25H01 | Pemberton | 50.35 N 122.83 W | radio | PRJNA979932 |
| AH25H02 | Pemberton | 50.35 N 122.83 W | radio | PRJNA979932 |
| AH25H03 | Pemberton | 50.41 N 122.89 W | radio | PRJNA979932 |
| AH26H01_S157_L001 | Pemberton | 50.26 N 122.87 W | radio | PRJNA979932 |
| AH26H02 | Pemberton | 50.26 N 122.87 W | radio | PRJNA979932 |
| AH26H03_S158_L001 | Pemberton | 50.26 N 122.87 W | radio | PRJNA1024534 |
| AH26H04 | Pemberton | 50.26 N 122.87 W | radio | PRJNA979932 |
| AH26H05_S159_L001 | Pemberton | 50.26 N 122.87 W | radio | PRJNA1024534 |
| AH26H06 | Pemberton | 50.26 N 122.87 W | radio | PRJNA979932 |
| AH26H07_S160_L001 | Pemberton | 50.26 N 122.87 W | radio | PRJNA1024534 |
| AH26H08 | Pemberton | 50.26 N 122.87 W | radio | PRJNA979932 |
| AH26H09 | Pemberton | 50.26 N 122.87 W | radio | PRJNA979932 |
| AH26H10 | Pemberton | 50.26 N 122.87 W | radio | PRJNA979932 |
| AH26H11 | Pemberton | 50.26 N 122.87 W | radio | PRJNA979932 |
| AH26H12_S161_L001 | Pemberton | 50.26 N 122.87 W | radio | PRJNA979932 |
| AH26H13 | Pemberton | 50.26 N 122.87 W | radio | PRJNA979932 |
| AH31H01 | Pemberton | 50.22 N 122.88 W | radio | PRJNA979932 |
| AH31H02 | Pemberton | 50.22 N 122.88 W | radio | PRJNA979932 |
| AH31H03 | Pemberton | 50.22 N 122.88 W | radio | PRJNA979932 |
| AH31H04 | Pemberton | 50.22 N 122.88 W | radio | PRJNA979932 |
| AI01H01 | Pemberton | 50.22 N 122.88 W | radio | PRJNA979932 |
| AI01H02 | Pemberton | 50.22 N 122.88 W | radio | PRJNA1024534 |
| AI01H03 | Pemberton | 50.22 N 122.88 W | radio | PRJNA979932 |
| AI01H04 | Pemberton | 50.22 N 122.88 W | radio | PRJNA979932 |
| AI01H05 | Pemberton | 50.22 N 122.88 W | radio | PRJNA979932 |
| AI01H06 | Pemberton | 50.22 N 122.88 W | radio | PRJNA979932 |
| AI01H07 | Pemberton | 50.22 N 122.88 W | radio | PRJNA979932 |
| AI01H08 | Pemberton | 50.22 N 122.88 W | radio | PRJNA979932 |
| AI01H09 | Pemberton | 50.22 N 122.88 W | radio | PRJNA1024534 |
| AI01H10 | Pemberton | 50.22 N 122.88 W | radio | PRJNA979932 |
| AI01H11 | Pemberton | 50.22 N 122.88 W | radio | PRJNA1024534 |
| AI01H12 | Pemberton | 50.22 N 122.88 W | radio | PRJNA979932 |
| AI01H13 | Pemberton | 50.22 N 122.88 W | radio | PRJNA979932 |
| AI01H14 | Pemberton | 50.22 N 122.88 W | radio | PRJNA979932 |
| AI01H15 | Pemberton | 50.22 N 122.88 W | radio | PRJNA979932 |
| AI01H16 | Pemberton | 50.22 N 122.88 W | radio | PRJNA979932 |
| AI01H17 | Pemberton | 50.22 N 122.88 W | radio | PRJNA979932 |
| AI02H01 | Pemberton | 50.22 N 122.88 W | radio | PRJNA979932 |
| AI02H03 | Pemberton | 50.22 N 122.88 W | radio | PRJNA979932 |
| AI02H04 | Pemberton | 50.22 N 122.88 W | radio | PRJNA979932 |
| AI02H05 | Pemberton | 50.22 N 122.88 W | radio | PRJNA979932 |
| AI02H06 | Pemberton | 50.22 N 122.88 W | radio | PRJNA979932 |
| AI02H07 | Pemberton | 50.22 N 122.88 W | radio | PRJNA979932 |
| AI02H08 | Pemberton | 50.22 N 122.88 W | radio | PRJNA979932 |
| AI02H10 | Pemberton | 50.22 N 122.88 W | radio | PRJNA979932 |
| BF23H01 | Alaska | 55.94 N 130.04 W | archival | PRJNA1024534 |
| BF23H02 | Alaska | 55.95 N 130.05 W | archival | PRJNA1024534 |
| BF24H01 | Alaska | 55.94 N 130.05 W | archival | PRJNA1024534 |
| BF24H02 | Alaska | 55.95 N 130.05 W | archival | PRJNA979932 |
| BF24H03 | Alaska | 55.96 N 130.06 W | archival | PRJNA1024534 |
| BF24H04 | Alaska | 55.96 N 130.06 W | archival | PRJNA1024534 |
| BF24H06 | Alaska | 55.94 N 130 W | archival | PRJNA1024534 |
| BF24H08 | Alaska | 55.94 N 129.99 W | archival | PRJNA979932 |
| BF25H01 | Alaska | 55.94 N 129.99 W | archival | PRJNA1024534 |
| BF25H02 | Alaska | 55.94 N 129.99 W | archival | PRJNA979932 |
| BF25H03 | Alaska | 55.94 N 130 W | archival | PRJNA979932 |
| BF25H04 | Alaska | 55.97 N 130.06 W | archival | PRJNA1024534 |
| BF26H01 | Alaska | 56.05 N 129.9 W | archival | PRJNA1024534 |
| BF26H02 | Alaska | 56.05 N 129.9 W | archival | PRJNA1024534 |
| BF26H03 | Alaska | 56.05 N 129.9 W | archival | PRJNA979932 |
| BF27H01 | Alaska | 56.05 N 129.9 W | archival | PRJNA1024534 |
| BF27H02 | Alaska | 56.05 N 129.9 W | archival | PRJNA1024534 |
| BF27H03 | Alaska | 56.04 N 129.9 W | archival | PRJNA1024534 |
| BF27H06 | Alaska | 56.04 N 129.9 W | archival | PRJNA979932 |
| BF27H07 | Alaska | 56.64 N 129.9 W | archival | PRJNA1024534 |
| BF28H01 | Alaska | 56.04 N 129.9 W | archival | PRJNA979932 |
| BF28H02 | Alaska | 56.04 N 129.9 W | archival | PRJNA1024534 |
| BF28H03 | Alaska | 56.04 N 129.9 W | archival | PRJNA979932 |
| BF28H04 | Alaska | 56.04 N 129.9 W | archival | PRJNA1024534 |
| BH28H03 | Pemberton | 50.22 N 122.88 W | radio | PRJNA979932 |
| BH28H04 | Pemberton | 50.22 N 122.88 W | radio | PRJNA979932 |
| BH28H05 | Pemberton | 50.22 N 122.88 W | radio | PRJNA979932 |
| BH28H06 | Pemberton | 50.22 N 122.88 W | radio | PRJNA979932 |
| BH28H07 | Pemberton | 50.26 N 122.87 W | radio | PRJNA1024534 |
| BH28H08 | Pemberton | 50.26 N 122.87 W | radio | PRJNA979932 |
| BH29H06 | Pemberton | 50.3 N 122.76 W | radio | PRJNA979932 |
| BH29H09 | Pemberton | 50.3 N 122.76 W | radio | PRJNA979932 |
| BH29H10 | Pemberton | 50.3 N 122.76 W | radio | PRJNA979932 |
| BH30H07 | Pemberton | 50.3 N 122.76 W | radio | PRJNA979932 |
| BH31H01 | Pemberton | 50.22 N 122.88 W | radio | PRJNA979932 |
| BH31H02_S173_L001 | Pemberton | 50.22 N 122.88 W | radio | PRJNA979932 |
| BH31H03 | Pemberton | 50.22 N 122.88 W | radio | PRJNA979932 |
| BH31H05 | Pemberton | 50.22 N 122.88 W | radio | PRJNA979932 |
| BH31H06_S174_L001 | Pemberton | 50.22 N 122.88 W | radio | PRJNA979932 |
| BH31H07 | Pemberton | 50.22 N 122.88 W | radio | PRJNA979932 |
| BH31H08 | Pemberton | 50.22 N 122.88 W | radio | PRJNA979932 |
| BH31H09 | Pemberton | 50.22 N 122.88 W | radio | PRJNA979932 |
| BH31H10_S175_L001 | Pemberton | 50.26 N 122.87 W | radio | PRJNA1024534 |
| BH31H11_S176_L001 | Pemberton | 50.3 N 122.76 W | radio | PRJNA1024534 |
| BH31H12 | Pemberton | 50.3 N 122.76 W | radio | PRJNA979932 |
| BH31H13_S177_L001 | Pemberton | 50.3 N 122.76 W | radio | PRJNA1024534 |
| BI01H01 | Pemberton | 50.22 N 122.89 W | radio | PRJNA979932 |
| BI01H02 | Pemberton | 50.22 N 122.89 W | radio | PRJNA979932 |
| BI01H03 | Pemberton | 50.22 N 122.88 W | radio | PRJNA979932 |
| BI01H04 | Pemberton | 50.22 N 122.88 W | radio | PRJNA979932 |
| BI01H05_S178_L001 | Pemberton | 50.22 N 122.88 W | radio | PRJNA979932 |
| BI01H06 | Pemberton | 50.22 N 122.88 W | radio | PRJNA979932 |
| BI01H07 | Pemberton | 50.22 N 122.88 W | radio | PRJNA979932 |
| BI01H08 | Pemberton | 50.22 N 122.88 W | radio | PRJNA979932 |
| BI01H09_S179_L001 | Pemberton | 50.22 N 122.88 W | radio | PRJNA1024534 |
| BI01H10 | Pemberton | 50.22 N 122.88 W | radio | PRJNA979932 |
| BI01H11 | Pemberton | 50.3 N 122.76 W | radio | PRJNA979932 |
| BI01H12 | Pemberton | 50.3 N 122.76 W | radio | PRJNA979932 |
| BI01H13 | Pemberton | 50.3 N 122.76 W | radio | PRJNA979932 |
| BI01H14 | Pemberton | 50.3 N 122.76 W | radio | PRJNA979932 |
| BI02H01 | Pemberton | 50.22 N 122.88 W | radio | PRJNA979932 |
| BI02H02 | Pemberton | 50.22 N 122.88 W | radio | PRJNA979932 |
| BI02H03 | Pemberton | 50.22 N 122.88 W | radio | PRJNA979932 |
| BI02H04 | Pemberton | 50.22 N 122.88 W | radio | PRJNA979932 |
| BI02H05 | Pemberton | 50.22 N 122.88 W | radio | PRJNA979932 |
| BI02H06 | Pemberton | 50.22 N 122.88 W | radio | PRJNA979932 |
| BI02H07_S180_L001 | Pemberton | 50.22 N 122.88 W | radio | PRJNA979932 |
| BI02H08 | Pemberton | 50.22 N 122.88 W | radio | PRJNA979932 |
| BI02H09 | Pemberton | 50.22 N 122.88 W | radio | PRJNA979932 |
| BI02H10 | Pemberton | 50.3 N 122.76 W | radio | PRJNA979932 |
| BI02H11 | Pemberton | 50.3 N 122.76 W | radio | PRJNA979932 |
| BI02H12 | Pemberton | 50.3 N 122.76 W | radio | PRJNA979932 |
| BI03H01 | Pemberton | 50.22 N 122.88 W | radio | PRJNA979932 |
| BI03H02 | Pemberton | 50.22 N 122.88 W | radio | PRJNA979932 |
| BI03H03 | Pemberton | 50.22 N 122.88 W | radio | PRJNA979932 |
| BI03H04 | Pemberton | 50.22 N 122.88 W | radio | PRJNA979932 |
| BI03H05_S181_L001 | Pemberton | 50.22 N 122.88 W | radio | PRJNA979932 |
| BI03H06 | Pemberton | 50.22 N 122.88 W | radio | PRJNA979932 |
| BI03H07 | Pemberton | 50.22 N 122.88 W | radio | PRJNA979932 |
| BI03H08 | Pemberton | 50.22 N 122.88 W | radio | PRJNA979932 |
| BI04H01 | Pemberton | 50.22 N 122.88 W | radio | PRJNA979932 |
| BI04H02_S182_L001 | Pemberton | 50.22 N 122.88 W | radio | PRJNA1024534 |
| BI04H03 | Pemberton | 50.22 N 122.88 W | radio | PRJNA979932 |
| BI04H04_S183_L001 | Pemberton | 50.22 N 122.88 W | radio | PRJNA1024534 |
| BI04H05 | Pemberton | 50.22 N 122.88 W | radio | PRJNA979932 |
| BI04H06_S184_L001 | Pemberton | 50.22 N 122.88 W | radio | PRJNA979932 |
| BI04H07 | Pemberton | 50.22 N 122.88 W | radio | PRJNA979932 |
| BI04H08 | Pemberton | 50.22 N 122.88 W | radio | PRJNA979932 |
| BI04H09 | Pemberton | 50.22 N 122.88 W | radio | PRJNA979932 |
| BI04H10 | Pemberton | 50.22 N 122.89 W | radio | PRJNA979932 |
| BI04H11 | Pemberton | 50.22 N 122.88 W | radio | PRJNA979932 |
| BI04H12 | Pemberton | 50.22 N 122.88 W | radio | PRJNA979932 |
| BI04H13 | Pemberton | 50.22 N 122.88 W | radio | PRJNA979932 |
| BI04H14 | Pemberton | 50.22 N 122.88 W | radio | PRJNA979932 |
| BI05H01 | Pemberton | 50.22 N 122.88 W | radio | PRJNA979932 |
| BI05H02 | Pemberton | 50.22 N 122.88 W | radio | PRJNA979932 |
| BI05H03_S185_L001 | Pemberton | 50.22 N 122.88 W | radio | PRJNA979932 |
| BI05H04 | Pemberton | 50.22 N 122.88 W | radio | PRJNA979932 |
| BI05H05_S186_L001 | Pemberton | 50.22 N 122.88 W | radio | PRJNA1024534 |
| BI05H06 | Pemberton | 50.22 N 122.88 W | radio | PRJNA979932 |
| BI05H07 | Pemberton | 50.22 N 122.88 W | radio | PRJNA979932 |
| BI05H10 | Pemberton | 50.26 N 122.87 W | radio | PRJNA979932 |
| BI05H11_S187_L001 | Pemberton | 50.26 N 122.87 W | radio | PRJNA979932 |
| BI05H12 | Pemberton | 50.26 N 122.87 W | radio | PRJNA979932 |
| BI05H13 | Pemberton | 50.3 N 122.76 W | radio | PRJNA979932 |
| BI06H01 | Pemberton | 50.26 N 122.87 W | radio | PRJNA979932 |
| BI06H02_S188_L001 | Pemberton | 50.26 N 122.87 W | radio | PRJNA1024534 |
| BI06H03 | Pemberton | 50.26 N 122.87 W | radio | PRJNA979932 |
| BI06H04 | Pemberton | 50.26 N 122.87 W | radio | PRJNA979932 |
| BI06H05 | Pemberton | 50.22 N 122.88 W | radio | PRJNA979932 |
| BI06H06 | Pemberton | 50.22 N 122.88 W | radio | PRJNA979932 |
| BI06H07 | Pemberton | 50.22 N 122.88 W | radio | PRJNA979932 |
| BI06H08 | Pemberton | 50.22 N 122.88 W | radio | PRJNA979932 |
| BI06H10 | Pemberton | 50.22 N 122.88 W | radio | PRJNA979932 |
| BI06H11_S189_L001 | Pemberton | 50.22 N 122.88 W | radio | PRJNA979932 |
| BI06H12 | Pemberton | 50.3 N 122.76 W | radio | PRJNA979932 |
| BI06H13_S190_L001 | Pemberton | 50.3 N 122.76 W | radio | PRJNA1024534 |
| BI06H14_S191_L001 | Pemberton | 50.3 N 122.76 W | radio | PRJNA1024534 |
| BI07H01_S192_L001 | Pemberton | 50.22 N 122.88 W | radio | PRJNA1024534 |
| BI07H02 | Pemberton | 50.22 N 122.88 W | radio | PRJNA979932 |
| BI07H03 | Pemberton | 50.22 N 122.88 W | radio | PRJNA979932 |
| BI07H04 | Pemberton | 50.22 N 122.88 W | radio | PRJNA979932 |
| bird_A | Washington | 47.28 N 121.08 W | archival | PRJNA979932 |
| bird_B | Washington | 47.28 N 121.08 W | archival | PRJNA1024534 |
| bird_C | Washington | 47.28 N 121.08 W | archival | PRJNA1024534 |
| bird_E | Washington | 47.29 N 121.09 W | archival | PRJNA1024534 |
| bird_F | Washington | 47.41 N 121.09 W | archival | PRJNA979932 |
| bird_G | Washington | 47.41 N 121.1 W | archival | PRJNA979932 |
| bird_H | Washington | 47.28 N 121.09 W | archival | PRJNA1024534 |
| bird_I | Washington | 47.36 N 121.11 W | archival | PRJNA1024534 |
| bird_J | Washington | 47.42 N 121.08 W | archival | PRJNA1024534 |
| bird_K | Washington | 47.42 N 121.08 W | archival | PRJNA1024534 |
| bird_M | Washington | 47.29 N 121.1 W | archival | PRJNA1024534 |
| bird_N | Washington | 47.35 N 121.1 W | archival | PRJNA1024534 |
| bird_O | Washington | 47.36 N 121.1 W | archival | PRJNA1024534 |
| bird_P | Washington | 47.36 N 121.1 W | archival | PRJNA1024534 |
| bird_Q | Washington | 47.36 N 121.1 W | archival | PRJNA1024534 |
| bird_S | Washington | 47.37 N 121.09 W | archival | PRJNA979932 |
| bird_T | Washington | 47.37 N 121.09 W | archival | PRJNA979932 |
| bird_U | Washington | 47.37 N 121.1 W | archival | PRJNA1024534 |
| bird_V | Washington | 47.38 N 121.1 W | archival | PRJNA1024534 |
| bird_W | Washington | 47.38 N 121.1 W | archival | PRJNA1024534 |
| bird_X | Washington | 47.41 N 121.11 W | archival | PRJNA1024534 |
| bird_Y | Washington | 47.41 N 121.11 W | archival | PRJNA1024534 |
| bird_Z | Washington | 47.19 N 121.01 W | archival | PRJNA1024534 |
| bird_Z_plus | Washington | 47.11 N 121.01 W | archival | PRJNA1024534 |
| CF18H03_S1_L001 | Hope | 49.28 N 121.15 W | archival | PRJNA979932 |
| CF19H02_S2_L001 | Hope | 49.28 N 121.15 W | archival | PRJNA1024534 |
| CF19H03_S3_L001 | Hope | 49.38 N 121.32 W | archival | PRJNA1024534 |
| CF19H04_S4_L001 | Hope | 49.23 N 121.19 W | archival | PRJNA1024534 |
| CF19H05_S5_L001 | Hope | 49.31 N 121.42 W | archival | PRJNA979932 |
| CF19H06_S6_L001 | Hope | 49.31 N 121.42 W | archival | PRJNA1024534 |
| CF19H07_S7_L001 | Hope | 49.32 N 121.42 W | archival | PRJNA1024534 |
| CF19H08_S8_L001 | Hope | 49.31 N 121.41 W | archival | PRJNA1024534 |
| CF19H09_S9_L001 | Hope | 49.31 N 121.41 W | archival | PRJNA1024534 |
| CF20H01_S10_L001 | Hope | 49.28 N 121.15 W | archival | PRJNA1024534 |
| CF20H03_S11_L001 | Hope | 49.48 N 121.25 W | archival | PRJNA1024534 |
| CF20H04_S12_L001 | Hope | 49.48 N 121.25 W | archival | PRJNA1024534 |
| CF20H05_S13_L001 | Hope | 49.32 N 121.42 W | archival | PRJNA1024534 |
| CF21H02_S15_L001 | Pemberton | 50.34 N 122.75 W | archival | PRJNA1024534 |
| CF21H03_S16_L001 | Pemberton | 50.34 N 122.75 W | archival | PRJNA979932 |
| CF21H04_S17_L001 | Pemberton | 49.32 N 121.42 W | archival | PRJNA979932 |
| CF21H05_S18_L001 | Pemberton | 50.33 N 122.75 W | archival | PRJNA979932 |
| CF22H01_S19_L001 | Pemberton | 50.33 N 122.74 W | archival | PRJNA1024534 |
| CF22H02_S20_L001 | Pemberton | 50.34 N 122.75 W | archival | PRJNA1024534 |
| CF22H03_S21_L001 | Pemberton | 50.21 N 122.88 W | archival | PRJNA1024534 |
| CF22H04_S22_L001 | Pemberton | 50.22 N 122.89 W | archival | PRJNA1024534 |
| CF22H05_S23_L001 | Pemberton | 50.22 N 122.88 W | archival | PRJNA1024534 |
| CF22H06_S24_L001 | Pemberton | 50.22 N 122.89 W | archival | PRJNA1024534 |
| CF23H01_S25_L001 | Pemberton | 50.21 N 122.88 W | archival | PRJNA1024534 |
| CF26H01_S26_L001 | Washington | 47.36 N 121.12 W | archival | PRJNA1024534 |
| CF27H01_S27_L001 | Washington | 47.32 N 121.1 W | archival | PRJNA1024534 |
| CF27H02_S28_L001 | Washington | 47.41 N 121.1 W | archival | PRJNA979932 |
| CF30H01_S29_L001 | Washington | 47.37 N 121.1 W | archival | PRJNA1024534 |
| CG01H01_S30_L001 | Washington | 47.38 N 121.1 W | archival | PRJNA1024534 |
| CG01H02_S31_L001 | Washington | 47.41 N 121.11 W | archival | PRJNA1024534 |
| CG01H03_S32_L001 | Washington | 47.38 N 121.1 W | archival | PRJNA1024534 |
| CG02H03_S33_L001 | Washington | 47.38 N 121.1 W | archival | PRJNA1024534 |
| CG05H01_S34_L001 | Washington | 47.42 N 121.08 W | archival | PRJNA1024534 |
| CH27H01_S35_L001 | Pemberton | 50.22 N 122.89 W | radio | PRJNA979932 |
| CH27H02_S36_L001 | Pemberton | 50.22 N 122.89 W | radio | PRJNA1024534 |
| CH27H03_S37_L001 | Pemberton | 50.22 N 122.89 W | radio | PRJNA979932 |
| CH27H04_S38_L001 | Pemberton | 50.22 N 122.89 W | radio | PRJNA979932 |
| CH28H01_S40_L001 | Pemberton | 50.26 N 122.87 W | radio | PRJNA979932 |
| CH28H02_S41_L001 | Pemberton | 50.26 N 122.87 W | radio | PRJNA1024534 |
| CH28H03_S42_L001 | Pemberton | 50.22 N 122.89 W | radio | PRJNA979932 |
| CH28H04_S43_L001 | Pemberton | 50.22 N 122.89 W | radio | PRJNA979932 |
| CH28H06_S44_L001 | Pemberton | 50.22 N 122.89 W | radio | PRJNA979932 |
| CH28H07_S45_L001 | Pemberton | 50.22 N 122.89 W | radio | PRJNA979932 |
| CH28H08_S46_L001 | Pemberton | 50.22 N 122.89 W | radio | PRJNA979932 |
| CH28H09_S47_L001 | Pemberton | 50.22 N 122.89 W | radio | PRJNA979932 |
| CH28H10_S48_L001 | Pemberton | 50.22 N 122.89 W | radio | PRJNA1024534 |
| CH28H11_S49_L001 | Pemberton | 50.22 N 122.89 W | radio | PRJNA979932 |
| CH28H12_S50_L001 | Pemberton | 50.22 N 122.89 W | radio | PRJNA979932 |
| CH29H01_S51_L001 | Pemberton | 50.26 N 122.87 W | radio | PRJNA979932 |
| CH29H02_S52_L001 | Pemberton | 50.26 N 122.87 W | radio | PRJNA1024534 |
| CH29H03_S53_L001 | Pemberton | 50.22 N 122.89 W | radio | PRJNA979932 |
| CH29H05_S55_L001 | Pemberton | 50.22 N 122.89 W | radio | PRJNA979932 |
| CH29H06_S56_L001 | Pemberton | 50.22 N 122.89 W | radio | PRJNA979932 |
| CH29H07_S57_L001 | Pemberton | 50.22 N 122.89 W | radio | PRJNA979932 |
| CH29H08_S58_L001 | Pemberton | 50.22 N 122.89 W | radio | PRJNA979932 |
| CH29H09_S59_L001 | Pemberton | 50.22 N 122.89 W | radio | PRJNA979932 |
| CH29H10_S60_L001 | Pemberton | 50.22 N 122.89 W | radio | PRJNA979932 |
| CH29H14_S62_L001 | Pemberton | 50.3 N 122.76 W | radio | PRJNA979932 |
| CH29H15_S63_L001 | Pemberton | 50.3 N 122.76 W | radio | PRJNA979932 |
| CH30H01_S64_L001 | Pemberton | 50.26 N 122.87 W | radio | PRJNA979932 |
| CH30H02_S65_L001 | Pemberton | 50.22 N 122.89 W | radio | PRJNA979932 |
| CH30H03_S66_L001 | Pemberton | 50.22 N 122.89 W | radio | PRJNA979932 |
| CH30H04_S67_L001 | Pemberton | 50.22 N 122.89 W | radio | PRJNA979932 |
| CH30H05_S68_L001 | Pemberton | 50.22 N 122.89 W | radio | PRJNA979932 |
| CH30H06_S69_L001 | Pemberton | 50.22 N 122.89 W | radio | PRJNA979932 |
| CH30H07_S70_L001 | Pemberton | 50.22 N 122.89 W | radio | PRJNA979932 |
| CH30H08_S71_L001 | Pemberton | 50.22 N 122.89 W | radio | PRJNA979932 |
| CH30H09_S72_L001 | Pemberton | 50.22 N 122.89 W | radio | PRJNA1024534 |
| CH30H10_S73_L001 | Pemberton | 50.22 N 122.89 W | radio | PRJNA979932 |
| CH30H13_S74_L001 | Pemberton | 50.3 N 122.76 W | radio | PRJNA979932 |
| CH31H01_S75_L001 | Pemberton | 50.22 N 122.89 W | radio | PRJNA979932 |
| CH31H02_S76_L001 | Pemberton | 50.26 N 122.87 W | radio | PRJNA979932 |
| CH31H03_S77_L001 | Pemberton | 50.26 N 122.87 W | radio | PRJNA979932 |
| CH31H04_S78_L001 | Pemberton | 50.26 N 122.87 W | radio | PRJNA979932 |
| CH31H05_S79_L001 | Pemberton | 50.26 N 122.87 W | radio | PRJNA979932 |
| CH31H06_S80_L001 | Pemberton | 50.22 N 122.89 W | radio | PRJNA979932 |
| CH31H07_S81_L001 | Pemberton | 50.22 N 122.89 W | radio | PRJNA979932 |
| CH31H10_S82_L001 | Pemberton | 50.22 N 122.89 W | radio | PRJNA979932 |
| CI03H01_S83_L001 | Pemberton | 50.22 N 122.89 W | radio | PRJNA979932 |
| CI03H02_S84_L001 | Pemberton | 50.22 N 122.89 W | radio | PRJNA979932 |
| CI03H03_S85_L001 | Pemberton | 50.22 N 122.89 W | radio | PRJNA979932 |
| CI03H04_S86_L001 | Pemberton | 50.22 N 122.89 W | radio | PRJNA1024534 |
| CI03H05_S87_L001 | Pemberton | 50.22 N 122.89 W | radio | PRJNA979932 |
| CI03H06_S88_L001 | Pemberton | 50.22 N 122.89 W | radio | PRJNA1024534 |
| CI04H01_S89_L001 | Pemberton | 50.22 N 122.89 W | radio | PRJNA979932 |
| CI04H02_S90_L001 | Pemberton | 50.22 N 122.89 W | radio | PRJNA979932 |
| CI04H03_S91_L001 | Pemberton | 50.22 N 122.89 W | radio | PRJNA979932 |
| CI04H04_S92_L001 | Pemberton | 50.22 N 122.89 W | radio | PRJNA979932 |
| CI04H05_S93_L001 | Pemberton | 50.22 N 122.89 W | radio | PRJNA979932 |
| CI04H06_S94_L001 | Pemberton | 50.22 N 122.89 W | radio | PRJNA979932 |
| CI04H07_S95_L001 | Pemberton | 50.22 N 122.89 W | radio | PRJNA979932 |
| CI04H08_S96_L001 | Pemberton | 50.22 N 122.89 W | radio | PRJNA979932 |
| CI04H09_S97_L001 | Pemberton | 50.22 N 122.89 W | radio | PRJNA979932 |
| CI05H01_S98_L001 | Pemberton | 50.26 N 122.87 W | radio | PRJNA979932 |
| CI05H02_S99_L001 | Pemberton | 50.26 N 122.87 W | radio | PRJNA979932 |
| CI05H03_S100_L001 | Pemberton | 50.22 N 122.89 W | radio | PRJNA979932 |
| CI05H04_S101_L001 | Pemberton | 50.22 N 122.89 W | radio | PRJNA979932 |
| CI05H06_S103_L001 | Pemberton | 50.26 N 122.87 W | radio | PRJNA979932 |
| CI06H01_S104_L001 | Pemberton | 50.22 N 122.89 W | radio | PRJNA979932 |
| CI06H02_S105_L001 | Pemberton | 50.26 N 122.87 W | radio | PRJNA979932 |
| CI06H03_S106_L001 | Pemberton | 50.22 N 122.89 W | radio | PRJNA979932 |
| CI06H04_S107_L001 | Pemberton | 50.22 N 122.89 W | radio | PRJNA979932 |
| CI06H05_S108_L001 | Pemberton | 50.22 N 122.89 W | radio | PRJNA1024534 |
| CI06H06_S109_L001 | Pemberton | 50.22 N 122.89 W | radio | PRJNA979932 |
| CI06H07_S110_L001 | Pemberton | 50.22 N 122.89 W | radio | PRJNA979932 |
| CI06H08_S111_L001 | Pemberton | 50.22 N 122.89 W | radio | PRJNA1024534 |
| CI06H09_S112_L001 | Pemberton | 50.22 N 122.89 W | radio | PRJNA979932 |
| CI06H10_S113_L001 | Pemberton | 50.22 N 122.89 W | radio | PRJNA979932 |
| CI06H11_S114_L001 | Pemberton | 50.22 N 122.89 W | radio | PRJNA1024534 |
| CI07H01_S115_L001 | Pemberton | 50.22 N 122.89 W | radio | PRJNA979932 |
| CI07H02_S116_L001 | Pemberton | 50.22 N 122.89 W | radio | PRJNA979932 |
| CI07H03_S117_L001 | Pemberton | 50.22 N 122.89 W | radio | PRJNA979932 |
| CI07H04_S118_L001 | Pemberton | 50.22 N 122.89 W | radio | PRJNA1024534 |
| CI09H01_S119_L001 | Pemberton | 50.22 N 122.89 W | radio | PRJNA979932 |
| CI09H02_S120_L001 | Pemberton | 50.22 N 122.89 W | radio | PRJNA979932 |
| CI09H03_S121_L001 | Pemberton | 50.22 N 122.89 W | radio | PRJNA979932 |
| CI09H04_S122_L001 | Pemberton | 50.22 N 122.89 W | radio | PRJNA979932 |
| CI09H05_S123_L001 | Pemberton | 50.22 N 122.89 W | radio | PRJNA979932 |
| CI09H06_S124_L001 | Pemberton | 50.22 N 122.89 W | radio | PRJNA979932 |
| CI09H07_S125_L001 | Pemberton | 50.22 N 122.89 W | radio | PRJNA979932 |
| CI09H08_S126_L001 | Pemberton | 50.22 N 122.89 W | radio | PRJNA979932 |
| DE28K03 | Pemberton | 50.37 N 122.86 W | archival | PRJNA1024534 |
| DE28K04 | Pemberton | 50.37 N 122.86 W | archival | PRJNA1024534 |
| DF25H01 | Pemberton | 50.34 N 122.74 W | archival | PRJNA1024534 |
| DF25H02 | Pemberton | 50.22 N 122.88 W | archival | PRJNA1024534 |
| DF25H03 | Pemberton | 50.22 N 122.88 W | archival | PRJNA1024534 |
| DF26H01 | Pemberton | 50.34 N 122.74 W | archival | PRJNA1024534 |
| DF26H02 | Pemberton | 50.34 N 122.74 W | archival | PRJNA1024534 |
| DF26H03 | Pemberton | 50.34 N 122.74 W | archival | PRJNA1024534 |
| DF27H01 | Pemberton | 50.34 N 122.74 W | archival | PRJNA1024534 |
| DF27H02 | Pemberton | 50.34 N 122.74 W | archival | PRJNA1024534 |
| DF27H03 | Pemberton | 50.34 N 122.74 W | archival | PRJNA1024534 |
| DF27H05 | Pemberton | 50.33 N 122.75 W | archival | PRJNA1024534 |
| DF27H06 | Pemberton | 50.33 N 122.75 W | archival | PRJNA1024534 |
| DF27H07 | Pemberton | 50.22 N 122.89 W | archival | PRJNA1024534 |
| DF27H08 | Pemberton | 50.22 N 122.89 W | archival | PRJNA1024534 |
| DF28H01 | Pemberton | 50.22 N 122.88 W | archival | PRJNA1024534 |
| DF28H02 | Pemberton | 50.22 N 122.89 W | archival | PRJNA1024534 |
| DF28H05 | Pemberton | 50.3 N 122.76 W | archival | PRJNA1024534 |
| DF28H07 | Pemberton | 50.3 N 122.76 W | archival | PRJNA1024534 |
| DF29H03 | Pemberton | 50.33 N 122.74 W | archival | PRJNA1024534 |
| DF29H04 | Pemberton | 50.34 N 122.75 W | archival | PRJNA1024534 |
| DF29H05 | Pemberton | 50.33 N 122.74 W | archival | PRJNA1024534 |
| DF29H06 | Pemberton | 50.34 N 122.74 W | archival | PRJNA1024534 |
| DF29K01 | Pemberton | 50.36 N 122.73 W | archival | PRJNA1024534 |
| DF29K02 | Pemberton | 50.34 N 122.76 W | archival | PRJNA1024534 |
| DF30H03 | Pemberton | 50.34 N 122.74 W | archival | PRJNA1024534 |
| DF30H04 | Pemberton | 50.41 N 122.89 W | archival | PRJNA1024534 |
| DG01H01 | Hope | 49.46 N 121.25 W | archival | PRJNA1024534 |
| DG01H02 | Hope | 49.46 N 121.25 W | archival | PRJNA1024534 |
| DG01H04 | Hope | 49.46 N 121.25 W | archival | PRJNA1024534 |
| DG01H05 | Hope | 49.47 N 121.25 W | archival | PRJNA1024534 |
| DG01H07 | Hope | 49.47 N 121.25 W | archival | PRJNA1024534 |
| DG01H08 | Hope | 49.47 N 121.25 W | archival | PRJNA1024534 |
| DG02H01 | Hope | 49.47 N 121.25 W | archival | PRJNA1024534 |
| DG02H02 | Hope | 49.48 N 121.25 W | archival | PRJNA1024534 |
| DG02H03 | Hope | 49.48 N 121.25 W | archival | PRJNA1024534 |
| DG02H04 | Hope | 49.47 N 121.25 W | archival | PRJNA1024534 |
| DG02H05 | Hope | 49.48 N 121.25 W | archival | PRJNA1024534 |
| DG02H06 | Hope | 49.48 N 121.25 W | archival | PRJNA1024534 |
| DG02H07 | Hope | 49.48 N 121.25 W | archival | PRJNA1024534 |
| DG02H08 | Hope | 49.48 N 121.25 W | archival | PRJNA1024534 |
| DG03H01 | Hope | 49.32 N 121.42 W | archival | PRJNA1024534 |
| DG03H02 | Hope | 49.31 N 121.42 W | archival | PRJNA1024534 |
| DG03H03 | Hope | 49.31 N 121.42 W | archival | PRJNA1024534 |
| DG03H04 | Hope | 49.32 N 121.42 W | archival | PRJNA1024534 |
| DG03H05 | Hope | 49.31 N 121.42 W | archival | PRJNA1024534 |
| DG03H06 | Hope | 49.32 N 121.42 W | archival | PRJNA1024534 |
| DG03H07 | Hope | 49.38 N 121.32 W | archival | PRJNA1024534 |
| DG03H08 | Hope | 49.38 N 121.32 W | archival | PRJNA1024534 |
| DG03H09 | Hope | 49.38 N 121.32 W | archival | PRJNA1024534 |
| DG04H01 | Hope | 49.32 N 121.42 W | archival | PRJNA1024534 |
| DG04H02 | Hope | 49.32 N 121.42 W | archival | PRJNA1024534 |
| DG04H03 | Hope | 49.32 N 121.42 W | archival | PRJNA1024534 |
| DG04H04 | Hope | 49.31 N 121.41 W | archival | PRJNA1024534 |
| DH27H01 | Pemberton | 50.22 N 122.89 W | radio | PRJNA1024534 |
| DH27H03 | Pemberton | 50.22 N 122.89 W | radio | PRJNA1024534 |
| DH27H04 | Pemberton | 50.22 N 122.89 W | radio | PRJNA1024534 |
| DH27H05 | Pemberton | 50.22 N 122.89 W | radio | PRJNA1024534 |
| DH27H07 | Pemberton | 50.22 N 122.89 W | radio | PRJNA1024534 |
| DH27H08 | Pemberton | 50.22 N 122.89 W | radio | PRJNA1024534 |
| DH27H09 | Pemberton | 50.22 N 122.89 W | radio | PRJNA1024534 |
| DH27H10 | Pemberton | 50.22 N 122.89 W | radio | PRJNA1024534 |
| DH27H11 | Pemberton | 50.22 N 122.89 W | radio | PRJNA1024534 |
| DH27H12 | Pemberton | 50.22 N 122.89 W | radio | PRJNA1024534 |
| DH27H13 | Pemberton | 50.22 N 122.89 W | radio | PRJNA1024534 |
| DH28H01 | Pemberton | 50.26 N 122.87 W | radio | PRJNA1024534 |
| DH28H02 | Pemberton | 50.26 N 122.87 W | radio | PRJNA1024534 |
| DH28H03 | Pemberton | 50.26 N 122.87 W | radio | PRJNA1024534 |
| DH28H04 | Pemberton | 50.22 N 122.89 W | radio | PRJNA1024534 |
| DH28H05 | Pemberton | 50.22 N 122.89 W | radio | PRJNA1024534 |
| DH28H06 | Pemberton | 50.22 N 122.89 W | radio | PRJNA1024534 |
| DH28H07 | Pemberton | 50.22 N 122.89 W | radio | PRJNA1024534 |
| DH28H08 | Pemberton | 50.22 N 122.89 W | radio | PRJNA1024534 |
| DH28H09 | Pemberton | 50.22 N 122.89 W | radio | PRJNA1024534 |
| DH28H10 | Pemberton | 50.22 N 122.89 W | radio | PRJNA1024534 |
| DH28H11 | Pemberton | 50.22 N 122.89 W | radio | PRJNA1024534 |
| DH28H12 | Pemberton | 50.22 N 122.89 W | radio | PRJNA1024534 |
| DH28H13 | Pemberton | 50.22 N 122.89 W | radio | PRJNA1024534 |
| DH28H14 | Pemberton | 50.22 N 122.89 W | radio | PRJNA1024534 |
| DH28H16 | Pemberton | 50.3 N 122.76 W | radio | PRJNA1024534 |
| DH28H17 | Pemberton | 50.3 N 122.76 W | radio | PRJNA1024534 |
| DH28H18 | Pemberton | 50.3 N 122.76 W | radio | PRJNA1024534 |
| DH28H19 | Pemberton | 50.3 N 122.76 W | radio | PRJNA1024534 |
| DH29H01 | Pemberton | 50.26 N 122.87 W | radio | PRJNA1024534 |
| DH29H02 | Pemberton | 50.22 N 122.89 W | radio | PRJNA1024534 |
| DH29H03 | Pemberton | 50.22 N 122.89 W | radio | PRJNA1024534 |
| DH29H04 | Pemberton | 50.22 N 122.89 W | radio | PRJNA1024534 |
| DH29H05 | Pemberton | 50.22 N 122.89 W | radio | PRJNA1024534 |
| DH29H06 | Pemberton | 50.26 N 122.87 W | radio | PRJNA1024534 |
| DH29H07 | Pemberton | 50.22 N 122.89 W | radio | PRJNA1024534 |
| DH29H08 | Pemberton | 50.22 N 122.89 W | radio | PRJNA1024534 |
| DH29H10 | Pemberton | 50.22 N 122.89 W | radio | PRJNA1024534 |
| DH29H11 | Pemberton | 50.22 N 122.89 W | radio | PRJNA1024534 |
| DH29H12 | Pemberton | 50.22 N 122.89 W | radio | PRJNA1024534 |
| DH29H13 | Pemberton | 50.3 N 122.76 W | radio | PRJNA1024534 |
| DH29H14 | Pemberton | 50.3 N 122.76 W | radio | PRJNA1024534 |
| DH29H15 | Pemberton | 50.3 N 122.76 W | radio | PRJNA1024534 |
| DH29H16 | Pemberton | 50.3 N 122.76 W | radio | PRJNA1024534 |
| DH30H01 | Pemberton | 50.22 N 122.89 W | radio | PRJNA1024534 |
| DH30H02 | Pemberton | 50.22 N 122.89 W | radio | PRJNA1024534 |
| DH30H03 | Pemberton | 50.22 N 122.89 W | radio | PRJNA1024534 |
| DH30H04 | Pemberton | 50.22 N 122.89 W | radio | PRJNA1024534 |
| DH30H05 | Pemberton | 50.22 N 122.89 W | radio | PRJNA1024534 |
| DH30H06 | Pemberton | 50.22 N 122.89 W | radio | PRJNA1024534 |
| DH30H11 | Pemberton | 50.26 N 122.87 W | radio | PRJNA1024534 |
| DH30H13 | Pemberton | 50.22 N 122.89 W | radio | PRJNA1024534 |
| DH30H14 | Pemberton | 50.22 N 122.89 W | radio | PRJNA1024534 |
| DH30H15 | Pemberton | 50.22 N 122.89 W | radio | PRJNA1024534 |
| DH30H16 | Pemberton | 50.3 N 122.76 W | radio | PRJNA1024534 |
| DH30H17 | Pemberton | 50.3 N 122.76 W | radio | PRJNA1024534 |
| DH31H02 | Pemberton | 50.26 N 122.87 W | radio | PRJNA1024534 |
| DH31H03 | Pemberton | 50.22 N 122.89 W | radio | PRJNA1024534 |
| DH31H04 | Pemberton | 50.22 N 122.89 W | radio | PRJNA1024534 |
| DH31H05 | Pemberton | 50.22 N 122.89 W | radio | PRJNA1024534 |
| DH31H06 | Pemberton | 50.22 N 122.89 W | radio | PRJNA1024534 |
| DH31H07 | Pemberton | 50.22 N 122.89 W | radio | PRJNA1024534 |
| DH31H08 | Pemberton | 50.22 N 122.89 W | radio | PRJNA1024534 |
| DH31H09 | Pemberton | 50.22 N 122.89 W | radio | PRJNA1024534 |
| DH31H10 | Pemberton | 50.22 N 122.89 W | radio | PRJNA1024534 |
| DH31H11 | Pemberton | 50.22 N 122.89 W | radio | PRJNA1024534 |
| DH31H12 | Pemberton | 50.22 N 122.89 W | radio | PRJNA1024534 |
| DH31H13 | Pemberton | 50.22 N 122.89 W | radio | PRJNA1024534 |
| DH31H15 | Pemberton | 50.22 N 122.89 W | radio | PRJNA1024534 |
| DH31H16 | Pemberton | 50.22 N 122.89 W | radio | PRJNA1024534 |
| DH31H17 | Pemberton | 50.3 N 122.76 W | radio | PRJNA1024534 |
| DI01H01 | Pemberton | 50.26 N 122.87 W | radio | PRJNA1024534 |
| DI01H02 | Pemberton | 50.26 N 122.87 W | radio | PRJNA1024534 |
| DI01H03 | Pemberton | 50.26 N 122.87 W | radio | PRJNA1024534 |
| DI01H04 | Pemberton | 50.26 N 122.87 W | radio | PRJNA1024534 |
| DI01H05 | Pemberton | 50.26 N 122.87 W | radio | PRJNA1024534 |
| DI01H06 | Pemberton | 50.26 N 122.87 W | radio | PRJNA1024534 |
| DI01H07 | Pemberton | 50.22 N 122.89 W | radio | PRJNA1024534 |
| DI01H08 | Pemberton | 50.22 N 122.89 W | radio | PRJNA1024534 |
| DI01H09 | Pemberton | 50.22 N 122.89 W | radio | PRJNA1024534 |
| DI01H10 | Pemberton | 50.22 N 122.89 W | radio | PRJNA1024534 |
| DI01H11 | Pemberton | 50.22 N 122.89 W | radio | PRJNA1024534 |
| DI01H12 | Pemberton | 50.22 N 122.89 W | radio | PRJNA1024534 |
| DI01H14 | Pemberton | 50.22 N 122.89 W | radio | PRJNA1024534 |
| DI01H15 | Pemberton | 50.22 N 122.89 W | radio | PRJNA1024534 |
| DI02H01 | Pemberton | 50.22 N 122.89 W | radio | PRJNA1024534 |
| DI02H02 | Pemberton | 50.22 N 122.89 W | radio | PRJNA1024534 |
| DI02H03 | Pemberton | 50.22 N 122.89 W | radio | PRJNA1024534 |
| DI02H05 | Pemberton | 50.22 N 122.89 W | radio | PRJNA1024534 |
| DI02H06 | Pemberton | 50.22 N 122.89 W | radio | PRJNA1024534 |
| DI02H07 | Pemberton | 50.22 N 122.89 W | radio | PRJNA1024534 |
| DI02H08 | Pemberton | 50.22 N 122.89 W | radio | PRJNA1024534 |
| DI02H09 | Pemberton | 50.22 N 122.89 W | radio | PRJNA1024534 |
| DI02H10 | Pemberton | 50.22 N 122.89 W | radio | PRJNA1024534 |
| DI03H01 | Pemberton | 50.26 N 122.87 W | radio | PRJNA1024534 |
| DI03H02 | Pemberton | 50.26 N 122.87 W | radio | PRJNA1024534 |
| DI03H03 | Pemberton | 50.26 N 122.87 W | radio | PRJNA1024534 |
| DI03H04 | Pemberton | 50.26 N 122.87 W | radio | PRJNA1024534 |
| DI03H05 | Pemberton | 50.22 N 122.89 W | radio | PRJNA1024534 |
| DI03H06 | Pemberton | 50.22 N 122.89 W | radio | PRJNA1024534 |
| DI03H07 | Pemberton | 50.22 N 122.89 W | radio | PRJNA1024534 |
| DI03H08 | Pemberton | 50.22 N 122.89 W | radio | PRJNA1024534 |
| DI03H09 | Pemberton | 50.22 N 122.89 W | radio | PRJNA1024534 |
| DI03H10 | Pemberton | 50.22 N 122.89 W | radio | PRJNA1024534 |
| DI03H11 | Pemberton | 50.22 N 122.89 W | radio | PRJNA1024534 |
| DI03H12 | Pemberton | 50.22 N 122.89 W | radio | PRJNA1024534 |
| DI04H01 | Pemberton | 50.26 N 122.87 W | radio | PRJNA1024534 |
| DI04H02 | Pemberton | 50.22 N 122.89 W | radio | PRJNA1024534 |
| DI04H03 | Pemberton | 50.22 N 122.89 W | radio | PRJNA1024534 |
| DI04H04 | Pemberton | 50.22 N 122.89 W | radio | PRJNA1024534 |
| DI04H05 | Pemberton | 50.22 N 122.89 W | radio | PRJNA1024534 |
| DI04H07 | Pemberton | 50.22 N 122.89 W | radio | PRJNA1024534 |
| JF05K04 | Pemberton | 50.22 N 122.89 W | archival | PRJNA1024534 |
| JF18K04 | Hope | 49.38 N 121.42 W | archival | PRJNA1024534 |
| JF18K05 | Hope | 49.38 N 121.42 W | archival | PRJNA1024534 |
| JF18K07 | Hope | 49.39 N 121.32 W | archival | PRJNA1024534 |
| JF27K01 | Pemberton | 50.34 N 122.75 W | archival | PRJNA1024534 |
| JG02K01 | Pemberton | 50.22 N 122.89 W | archival | PRJNA1024534 |
| KF11K02 | Hope | 49.47 N 121.25 W | archival | PRJNA1024534 |
| KF11K03 | Hope | 49.48 N 121.25 W | archival | PRJNA1024534 |
| KF11K04 | Hope | 49.48 N 121.25 W | archival | PRJNA1024534 |
| KF11K05 | Hope | 49.48 N 121.25 W | archival | PRJNA1024534 |
| KF12K01 | Hope | 49.48 N 121.25 W | archival | PRJNA1024534 |
| KF18K01 | Hope | 49.38 N 121.42 W | archival | PRJNA1024534 |
| KF18K03 | Hope | 49.38 N 121.42 W | archival | PRJNA1024534 |
| KF18K05 | Hope | 49.38 N 121.42 W | archival | PRJNA1024534 |
| KF19K01 | Hope | 49.47 N 121.25 W | archival | PRJNA1024534 |
| KF19K02 | Hope | 49.47 N 121.25 W | archival | PRJNA1024534 |
| KF19K04 | Pemberton | 50.32 N 122.74 W | archival | PRJNA1024534 |
| KF19K05 | Pemberton | 50.33 N 122.74 W | archival | PRJNA1024534 |
| KF20K01 | Pemberton | 50.33 N 122.74 W | archival | PRJNA1024534 |
| KF20K03 | Pemberton | 50.34 N 122.74 W | archival | PRJNA1024534 |
| KF20K04 | Pemberton | 50.34 N 122.74 W | archival | PRJNA1024534 |
| KF20K05 | Pemberton | 50.34 N 122.74 W | archival | PRJNA1024534 |
| KF20K06 | Pemberton | 50.33 N 122.74 W | archival | PRJNA1024534 |
| KF20K08 | Pemberton | 50.34 N 122.73 W | archival | PRJNA1024534 |
| KF21K01 | Pemberton | 50.34 N 122.75 W | archival | PRJNA1024534 |
| KF21K02 | Pemberton | 50.34 N 122.75 W | archival | PRJNA1024534 |
| KF21K04 | Pemberton | 50.3 N 122.76 W | archival | PRJNA1024534 |
| KF21K06 | Pemberton | 50.3 N 122.76 W | archival | PRJNA1024534 |
| KF21K07 | Pemberton | 50.3 N 122.76 W | archival | PRJNA1024534 |
| KF22K01 | Pemberton | 50.3 N 122.76 W | archival | PRJNA1024534 |
| KF22K02 | Pemberton | 50.3 N 122.76 W | archival | PRJNA1024534 |
| KF22K03 | Pemberton | 50.29 N 122.76 W | archival | PRJNA1024534 |
| KF22K04 | Pemberton | 50.29 N 122.76 W | archival | PRJNA1024534 |
| KF22K05 | Pemberton | 50.29 N 122.76 W | archival | PRJNA1024534 |
| KF22K06 | Pemberton | 50.29 N 122.76 W | archival | PRJNA1024534 |
| KF22K07 | Pemberton | 50.3 N 122.76 W | archival | PRJNA1024534 |
| KF23K01 | Pemberton | 50.22 N 122.88 W | archival | PRJNA1024534 |
| KF23K02 | Pemberton | 50.22 N 122.89 W | archival | PRJNA1024534 |
| KF23K03 | Pemberton | 50.22 N 122.88 W | archival | PRJNA1024534 |
| KF23K04 | Pemberton | 50.22 N 122.88 W | archival | PRJNA1024534 |
| KF23K09 | Pemberton | 50.22 N 122.88 W | archival | PRJNA1024534 |
| KF24K01 | Pemberton | 50.35 N 122.73 W | archival | PRJNA1024534 |
| KF24K02 | Pemberton | 50.35 N 122.73 W | archival | PRJNA1024534 |
| KF25K01 | Hope | 49.38 N 121.41 W | archival | PRJNA1024534 |
| LE21K01 | Hope | 49.48 N 121.25 W | archival | PRJNA979932 |
| LF04K01 | Pemberton | 50.34 N 122.74 W | archival | PRJNA979932 |
| LF06K01 | Pemberton | 50.22 N 122.89 W | archival | PRJNA979932 |
| LF09K01 | Hope | 49.38 N 121.41 W | archival | PRJNA979932 |
| LF11K01 | Hope | 49.38 N 121.32 W | archival | PRJNA979932 |
| LF11K02 | Hope | 49.39 N 121.32 W | archival | PRJNA979932 |
| LF12K01 | Hope | 49.47 N 121.25 W | archival | PRJNA979932 |
| LF14K01 | Pemberton | 50.22 N 122.88 W | archival | PRJNA1024534 |
| LF16K01 | Pemberton | 50.3 N 122.76 W | archival | PRJNA979932 |
| LF17K01 | Pemberton | 50.33 N 122.74 W | archival | PRJNA979932 |
| LF21K01 | Pemberton | 50.28 N 122.89 W | archival | PRJNA1024534 |
| LF22K01 | Pemberton | 50.28 N 122.89 W | archival | PRJNA1024534 |
| LF23K01 | Pemberton | 50.28 N 122.89 W | archival | PRJNA1024534 |
| LF23K02 | Pemberton | 50.28 N 122.89 W | archival | PRJNA1024534 |
| LF23K03 | Pemberton | 50.28 N 122.89 W | archival | PRJNA1024534 |
| LF23K05 | Pemberton | 50.28 N 122.89 W | archival | PRJNA1024534 |
| LF23K06 | Pemberton | 50.28 N 122.89 W | archival | PRJNA1024534 |
| LF23K07 | Pemberton | 50.28 N 122.89 W | archival | PRJNA1024534 |
| LF24K02 | Pemberton | 50.28 N 122.89 W | archival | PRJNA1024534 |
| LF24K03 | Pemberton | 50.28 N 122.89 W | archival | PRJNA1024534 |
| LF24K06 | Pemberton | 50.28 N 122.89 W | archival | PRJNA1024534 |
| LF24K08 | Pemberton | 50.28 N 122.89 W | archival | PRJNA1024534 |
| LF25K01 | Pemberton | 50.28 N 122.89 W | archival | PRJNA1024534 |
| LF26K01 | Hope | 49.36 N 121.35 W | archival | PRJNA1024534 |
| LF26K03 | Hope | 49.36 N 121.35 W | archival | PRJNA1024534 |
| LF26K04 | Hope | 49.36 N 121.35 W | archival | PRJNA1024534 |
| LF26K05 | Hope | 49.47 N 121.25 W | archival | PRJNA979932 |
| LF26K06 | Hope | 49.36 N 121.35 W | archival | PRJNA1024534 |
| LF26K07 | Hope | 49.36 N 121.35 W | archival | PRJNA1024534 |
| LF26K09 | Hope | 49.36 N 121.35 W | archival | PRJNA1024534 |
| LF27K01 | Hope | 49.36 N 121.35 W | archival | PRJNA1024534 |
| LF27K05 | Hope | 49.36 N 121.35 W | archival | PRJNA1024534 |
| LF27K06 | Hope | 49.36 N 121.35 W | archival | PRJNA1024534 |
| LF27K09 | Hope | 49.36 N 121.35 W | archival | PRJNA1024534 |
| LF27K10 | Hope | 49.36 N 121.35 W | archival | PRJNA1024534 |
| LF27K11 | Hope | 49.36 N 121.35 W | archival | PRJNA1024534 |
| MF03K01 | Hope | 49.36 N 121.35 W | archival | PRJNA979932 |
| MF06K01 | Pemberton | 50.28 N 122.89 W | archival | PRJNA979932 |
| MF08K01 | Pemberton | 50.28 N 122.89 W | archival | PRJNA979932 |
| MF09K01 | Pemberton | 50.28 N 122.89 W | archival | PRJNA979932 |
| MF12K01 | Pemberton | 50.28 N 122.89 W | archival | PRJNA979932 |
| MF12K02 | Pemberton | 50.28 N 122.89 W | archival | PRJNA979932 |
| MF12K03 | Hope | 49.36 N 121.35 W | archival | PRJNA979932 |
| MF14K01 | Hope | 49.36 N 121.35 W | archival | PRJNA979932 |
| MF15K01 | Hope | 49.36 N 121.35 W | archival | PRJNA979932 |
| MF16K01 | Hope | 49.36 N 121.35 W | archival | PRJNA979932 |
| MF19K01 | Pemberton | 50.28 N 122.89 W | archival | PRJNA979932 |
